# Aldehydic load as an objective imaging biomarker of mild traumatic brain injury

**DOI:** 10.1101/2024.04.16.589820

**Authors:** Alexia Kirby, Cian Ward, Nicholas D. Calvert, Ryan Daniel, Joseph Wai-Hin Leung, Ashwin Sharma, Mojmír Suchý, Cassandra Donatelli, Jing Wang, Emily Standen, Adam J. Shuhendler

**Author notes:** Corresponding author: Adam Shuhendler.

## Abstract

Concussion is a mild traumatic brain injury (mTBI) defined as complex neurological impairment induced by biomechanical forces without structural brain damage. There does not yet exist an objective diagnostic tool for concussion. Downstream injury from mTBI stems from oxidative damage leading to the production of neurotoxic aldehydes. A collagen-based 3D corticomimetic scaffold was developed affording an *in vitro* model of concussion, which confirmed increased aldehyde production in live neurons following impact. To evaluate total aldehyde levels *in vivo* following mTBI, a novel CEST-MRI contrast agent, ProxyNA_3_, has been implemented in a new model of closed-head, awake, single-impact concussion developed in aged and young mice with aldehyde dehydrogenase 2 (ALDH2) deficiency. Behavioural tests confirm deficits immediately after injury. ProxyNA_3_-MRI was performed before impact, and on days two- and seven- post-impact. MRI signal enhancement significantly increased at two days post-injury and decreased to baseline seven days post-injury in all mice. An increase in astrocyte activation at seven days post-injury confirms the onset of a neuroinflammatory response following aldehyde production in the brain. The data suggest that advanced age and ALDH2 deficiency contribute to increased aldehydic load following mTBI. Overall, ProxyNA_3_ was capable of mapping concussion-associated aldehydes, supporting its application as an objective diagnostic tool for concussion.

## Introduction

More than 50 million people worldwide experience a Traumatic Brain Injury (TBI) each year, with mild TBI (mTBI) accounting for up to 90% of those cases ^[1]^. The incidence of sports-related concussions has also been on the rise, with a more than 100% increase from 2001 to 2012 in North America ^[2]^. It is important to note that mTBI statistics are prone to diagnostic and selection biases, as concussions are underreported, underdiagnosed, and misdiagnosed ^[1, 3]^. Objective, unambiguous diagnosis is crucial for safe return to activities to avoid further exacerbation of symptoms and development of neurological deficits later in life ^[4]^.

Currently, the detection of intracranial injuries relies on head Computed Tomography (CT), which may result in unnecessary ionizing radiation in young individuals with mTBI ^[5]^. Studies show 85-99% of adults and 93-100% of children with mTBI have no intracerebral lesions, in which cases cranial CT scans could be avoided ^[6]^. Unlike more severe TBIs, concussions often do not present visible physical trauma or external signs of injury with a delayed onset of symptoms that vary widely ^[7]^. Currently, there is no single objective diagnostic tool for concussions, but instead diagnosis relies on a combination of medical history, physical examination, symptom evaluation, and, in some cases, neuroimaging to rule out more severe traumatic brain injury ^[8]^. A direct, imaging-based assessment of brain health after mTBI is an outstanding clinical need.

The pathophysiology of mTBIs derives from the neurometabolic cascade of concussion, propagating from irregular ionic flux due to lipid membrane mechanoporation, resulting in an energy crisis, altered intracellular redox state, and formation of reactive oxygen species ^[9]^. The induced oxidative stress promotes polyamine catabolism and lipid peroxidation, resulting in the production of a diverse array of biogenic aldehydes detectable in rodents, human plasma, and human cerebral spinal fluid within 72 hours post-brain injury ^[10]^. Scavenging aldehydes in the early hours following TBI can have neuroprotective effects ^[11]^, suggesting aldehydes are not only markers of injury, but also early propagators of pathogenesis.

While aldehydes play a significant role in the early stages of TBI neuropathy ^[12]^, it is not yet clear whether advanced age leads to worse clinical outcomes following head injury, as TBI studies often exclude older adults due to pre-existing neurological conditions ^[13]^. Preclinical evidence exists supporting an increase in oxidative damage, lowered antioxidant capacities, and increased tissue loss in aged rats compared to young rats following moderate TBI ^[12c]^. The redox stress hypothesis of aging proposes a shift from predominately anti-oxidant to more pro-oxidant redox state as age progresses ^[14]^, increasing aldehyde production as secondary messengers of pathology, amplifying injury ^[15]^. With the recent shift of increased incidence of TBI in the older population ^[16]^, there is an urgent need for more investigation into this under-researched topic ^[17]^. The progressive decline of aldehyde metabolism with advancing age is, in part, due to a reduced activity of aldehyde dehydrogenases (ALDH), a family of enzymes responsible for the elimination of endogenous aldehydes ^[18]^. Mitochondrial ALDH2 plays a major role in the detoxification of aldehydes derived from lipid peroxidation ^[19]^. Given the redox hypothesis of aging, and evidence supporting aldehydes as propagators of injury, we hypothesize that older individuals will have a greater neural aldehyde load following mTBI, which may lead to more severe outcomes.

While the neurometabolic cascade of concussion has been shown to result in the elevated production of aldehydes that can serve as a potential mTBI biomarker, detecting and imaging aldehydic load in live subjects remains a challenge, hindering their use as diagnostic tools. In this study, the generation of aldehydes following mechanical injury of neurons has been shown, and the diagnostic potential of aldehydic load as a biomarker for mTBI has been demonstrated. A novel chemical exchange saturation transfer (CEST) magnetic resonance imaging (MRI) contrast agent, 5-propargyloxy-*N*-aminoanthranilic acid (ProxyNA_3_), has been developed to map aldehydic load in a mouse model of mTBI, affording the use of pathology-associated aldehydes as imaging biomarkers for concussion.

## Methods

### Animals

All procedures for animal experimentation were conducted under Animal Use Protocol SCe-4019 approved by the IACUC at the University of Ottawa. Mice were housed under 12 hr light/dark cycle, and ambient temperature of 20-24°C and 45 to 65% humidity. All mice were provided access to food (Rodent Laboratory Chow) and water *ad libitum*. The ALDH2 knockout mice have a C57BL/6 background and were generated as previously described ^[20]^, and kindly provided by Dr. Brian Bennett (Queen’s University, Kingston, Canada). Heterozygous and knockout mice were generated by breeding heterozygotes, genotyping offspring with extracted DNA from ear punches or tail snips by PCR ^[21]^. Wild type mice were obtained from Charles River Laboratories (MA, USA).

### Statistical Analyses

Statistical analyses were performed using Prism (v. 9.5.0, GraphPad, Inc.). Two-group comparisons were performed by two-tail Student’s t-test. Comparisons across more than two groups was performed by one-way ANOVA followed by Tukey’s test for honestly significant differences. Normality was assumed where appropriate for all data sets. Prior to ANOVA, Levene’s test was used to confirm equal variance, and visual quantile-quantile plot analysis was used to confirm homoscedasticity.

### Corticomimetic scaffold

#### Donut Scaffold Exterior Preparation

All steps were performed in a biological safety cabinet. Stock glutaraldehyde solution is diluted to 0.75% in a final volume of 30μL using PBS (Gibco). Stock 1M NaOH solution was used to neutralize collagen solution. Fibricol Type I collagen (Advanced Biomatrix) was added to a glass vial containing a magnetic stir bar and then supplemented with 10X DMEM and glutaraldehyde on ice. Collagen hydrogel was neutralized to a pH of 7.6 using 1.0M NaOH and left on ice in the dark for 15 minutes before dispensing the gel in 500 μL aliquots into 20 mm diameter coverwell imaging chambers (Electron Microscopy Sciences). Dispensed gels were incubated at 37°C overnight in a HERAcell 150 incubator (Thermo Fisher) in the coverslips to solidify. Solidified scaffolds were removed from the coverslips, placed in a 6 well tissue culture plate (VWR) and covered in 20% glycine for 1 hour at 37°C, 5.0% CO_2_ to quench unreacted glutaraldehyde. Collagen gel exteriors were then washed with three volumes of 3mL PBS (Gibco), before being placed back in their original coverslips. Using a 3.0 mm diameter disposable biopsy punch (Miltex), the centers of each cross-linked collagen gel was removed. Punched collagen hydrogels were then covered in neurobasal media and incubated at 37°C while the second collagen matrix was being prepared.

#### Primary cell culture

Primary cortical precursors were obtained from E11-12 cortices dissected from CD-1 mice (Charles River Laboratories) as previously described^[22]^. Briefly, embryos were transferred to ice-cold Hanks’ balanced salt solution (HBSS) (Fisher Scientific), meninges were removed, and cerebral cortices were isolated from the brain. The cortical tissue was mechanically triturated with a plastic pipette and seeded directly into a 24-well plate (Thermo Fisher Scientific), pre-coated with 15% poly-L-ornithine (PLO) (Sigma-Aldrich) and 5% laminin (Thermo Fisher Scientific). The cortical precursors were cultured following a neurosphere assay protocol^[23]^ in a neurobasal medium (Thermo Fisher Scientific) containing 500µM GlutaMAX supplement (Thermo Fisher Scientific), 2% B27 supplement (Thermo Fisher Scientific), 1% penicillin/streptomycin (Thermo Fisher Scientific), 40ng/mL endothelial growth factor (EGF; Thermo Fisher Scientific), and 40 ng/ml FGF2 (PeproTech). The seeded 24 well plate was incubated at 37°C, 5.0% CO_2_ for six days, with Neurobasal media changed daily. When the cells formed neurospheres (approximately six days after seeding), they were trypsinized, pelleted by centrifugation (110 x g for 15 min) at room temperature and resuspended in the appropriate volume of neurobasal medium to give a concentration of 1.28× 10^6^ cells/mL. Cortical precursor resuspension was kept on ice until needed.

#### Donut Scaffold Interior Preparation

A second collagen matrix was prepared by diluting Fibricol Type I collagen (Advanced Biomatrix) in the appropriate volume of 0.01M HCl to give a collagen concentration of 2.0 mg/mL. Collagen solution was mixed using a magnetic stir bar and was supplemented with 10X DMEM and neutralized to a pH of 7.2-7.6 using 1.0 M NaOH. After neutralization, cortical precursor cell suspension was added to give a final cell concentration of 1,000,000 cells. Sterile water was added to the collagen solution to give a final volume of 625 μL and the solution was mixed gently with a magnetic stir bar. Cortical precursor-seeded collagen was left on ice in the dark for 15 minutes before aliquoting in 30 μL volumes to fill the centers of the donut exteriors. Donut scaffolds were incubated for 1.5 hours at 37°C, 5.0% CO_2_ before being removed from their coverslips, placed in a 6 well tissue culture plate (VWR) and covered in Neurobasal Medium (Gibco) supplemented with 2% B27, 1% Penicillin/Streptomycin (Gibco), GlutaMAX (Gibco), 40ng/mL FGF2 (PeproTech) and 40ng/mL EGF (Thermo-Fisher) and incubated at 37°C, 5.0% CO_2_ for four days before imaging or performing any further experiments.

#### Mechanical testing with UTM

Scaffolds or 6 mm punches of dissected fresh mouse cortical brain tissue were placed on a Fisherbrand double frosted microscope slide (Fisher Scientific) in the center of an adhesive, water-tight silicone gasket (Electron Microscopy Sciences). PBS was added to surround, but not submerge the scaffolds/brain tissue. The soaked material was then compressed 50% of its thickness five times using a UTM. The UTM recorded the thickness and standard force of the tested material. All material was tested in quintuplicate.

##### UTM settings

Speed at starting position is 100mm/min, with a pre-load force of 0.002N, and pre-load speed of 50mm/min. A cyclic loading measurement phase with 5 cycles was applied with a strain of 50% at a speed of 3%/s, and a hold time of 1s at the point of load application and load removal of the cycles. The upper force limit was set to 20N.

#### Compression Testing Data Analysis

The Young’s Modulus value was used to evaluate the stiffness of the scaffolds and cortical brain tissue. Young’s Modulus was determined by taking the slope of the linear portion of a stress vs strain graph of the first compression of the material. Strain was measured as a percentage of the gel’s thickness and stress was calculated as the applied force per unit of material area. Young’s Modulus data across varying materials was compared using a Kruskal-Wallis non-parametric test to determine significance between the multiple experimental groups. When a Kruskal-Wallis test identified significance between experimental groups, a Benjamini-Hochberg test was performed to give adjusted p-values of each experimental group. Using the adjusted p-values of the Benjamini-Hochberg test, a Dunn’s post-hoc test was used to find significance between experimental groups. All statistical analysis was performed on R.

#### Compression and confocal imaging

##### Live/dead staining

Mouse primary neuron cells isolated and seeded into scaffolds as previously described were washed with PBS and then incubated with 1 mL of PBS containing calcein-acetoxymethyl ester (calcein-AM, 2 µM) and ethidium homodimer-1 (EthD-1, 4 µM) for 30 min under cell culture conditions. The scaffolds were washed gently three times with PBS and mounted onto a polymer-bottom 24 well plate (Ibidi). Scaffolds were imaged using a Zeiss LSM 880 confocal microscope with excitation/emission (ex./em.) and multi-bandpass filters (MPB) for calcein-AM set to 495/515 nm MBP 488nm and EthD-1 set to 495/635 nm MBP 458/561nm, respectively. Tile-scanned (3x3) Z-stacked images were acquired on 20x objective within the depth range of coherent light. Live cells fluoresced green (calcein-AM) and dead cells fluoresced red (EthD-1). Images were presented as maximum intensity projections (MIPs) of the Z-stacks.

##### NeuO

Cells are incubated with 0.25µM NeuO in cell culture medium at 37°C for 1 hour. The imaging medium was then replaced with fresh culture medium and incubated at 37°C for 2 hours before imaging. Confocal imaging was performed using a 20X objective with ex/em set to 488/557nm with multi-bandpass filter (MPB) of 488nm.

##### 5MeONA_3_

Images of live cells within the scaffold were taken after incubation 5MeONA_3_ for aldehyde detection, before and after impact. The cell-seeded scaffold was placed in a well of a 12 well polymer-bottom plate and submerged in phenol-free media. Images were taken at 10X, 20X, or 64X objective at multiple depths of focus, with ex/em set to 405/420-520nm and MBP 405nm. After pre-impact imaging, the scaffold was removed from the well containing media and placed on a microscope slide and positioned under a custom-made 30mm tip for the impactor device. The depth was set to 0.5mm, velocity 4m/s and dwell time 0.3s. After impact, the gel was placed back into the well containing phenol red-free media to proceed with post-impact imaging.

### mTBI mouse model

Model development including optimization of impact parameters, behavioral tests, and DCE-MRI was performed on WT Balb/c mice.

#### Awake closed head impact

Unanesthetized mice were placed head-first into a conical restraint bag and held in a prone position by the base of the tail and hindlimbs (Fig. S3A). The head of the mouse was positioned under the tip (5mm diameter) of the impactor (Leica Impact One Electromagnetic Impactor). The mouse was held over a sheet of filter floss to allow free linear and rotational movement of the head and neck. The velocity, depth, and dwell time were set (see Table S1) and the impact was performed, with the tip making contact around the site of bregma. Immediately after impact, the mice were given a subcutaneous bolus of buprenorphrine (0.01mg/kg) and placed in the supine position until righting reflex was regained. Sham mice were placed in restraint bag and held under impactor for 30 seconds, then removed from the bag to simulate the stress associated with handling.

#### Neurological Severity Score (NSS)

The neurological severity score (NSS) is made up of a variety of tasks used to assess a rodent’s balance, normal behavior, cognition, and reflexes (Supporting Fig. S3B-H). The procedure was adapted from previous studies ^[24]^. Tasks were judged in a pass/fail format with zero points being awarded for completing the task and one point being awarded for failing to complete the task. As such, higher NSS is used to indicate greater deficits in the abilities mentioned above, with a maximum composite score of 5. For model development, mice performed the NSS immediately after impact, and at 1-, 4- and 24-hours post-impact. Note for the cohort of mice impacted at 2.35 m/s, the NSS was not performed immediately after impact. Each of the tasks incorporated in the NSS are outlined below in the order in which they were performed:

1. Hindlimb Reflex. A mouse was raised approximately 60 cm in the air by the tail (Supporting Fig. S3D). A point was awarded if the mouse failed to extend both forelimbs and hindlimbs, and instead crossed its limbs.
2. Startle Reflex. A mouse was placed on the flat surface of the lab bench and one researcher clapped over top of the mouse, while the other researcher observed for the presence of a normal startle reflex (Supporting Fig. S3E). A point was awarded if the mouse failed to display a proper startle reflex.
3. Escape Paradigm. A mouse was placed in the center of a cylinder with a diameter of 30 cm, a height of 20 cm and a small door (5 cm^2^) to allow the mouse to escape (Supporting Fig. S3F). A point was awarded if the mouse failed to successfully exit the door during a 3-minute period.
4. Balance Test. A mouse was placed in the center of a meter stick positioned on its side (width of 0.6 cm) that was being held by a researcher in between an empty cage and the rodent’s home cage (Fig. S3G). A point was awarded if the mouse failed to balance on the meter stick in a perched position for a period of 10 seconds.
5. Beam Walk Test. Using the same meter stick from task 4., the mouse was again placed in the center of the stick that was supported by an empty cage and the rodent’s home cage that was placed 60 cm away (Fig. S3H). This time the meter stick was in the flat position (width of 2.5 cm). If the mouse failed to traverse 30 cm in either direction on the stick during a 3-minute period, a point was awarded.

#### Grid walk

Our modified grid walk test was used as an additional measure for acute deficits in motor coordination, as well as a measure of activity level^[25]^. The mouse was placed in the center of a 21 cm^2^ grid (1/2-inch hole size) elevated 30 cm in the air (Fig. S3B-H). A video camera was positioned at a 45-degree angle underneath the grid for visualization. Mice were recorded for a 3-minute period 24 hours before impact and immediately after the NSS was performed post-concussive impact. Analysis was performed by a blinded researcher watching the videos frame by frame. The total number of forelimb strides were recorded, and the total number of forelimb foot faults were recorded. A foot fault was defined as anytime the mouse misplaced one of its forelimbs allowing it to pass directly through the grid (Fig. S3B *vs*. C). The activity level of the mouse was also recorded during the 3-minute period. Being active was defined when any part of the mouse was moving, which included exploratory behavior, such as just moving the head. The percentage of forelimb foot faults was calculated by dividing the number of forelimb foot faults by the total number of forelimb steps. The percentage of time active was calculated by dividing the total time spent active by the 3-minute recording period.

### Probe preparation

5MeONA_3_^[26]^was synthesized as previously described; spectral characterization match that previously reported.

ProxyNA_3_ was prepared from 5-*O*-propargyl-*N*-Boc-anthranilic acid previously prepared by us ^[27]^ by modification of an existing literature protocol ^[28]^. A one pot reaction involving Boc deprotection and a diazotization/reduction cascade resulted in ProxyNA_3_ ·2 HCl (Scheme 1).

**Scheme 1.**
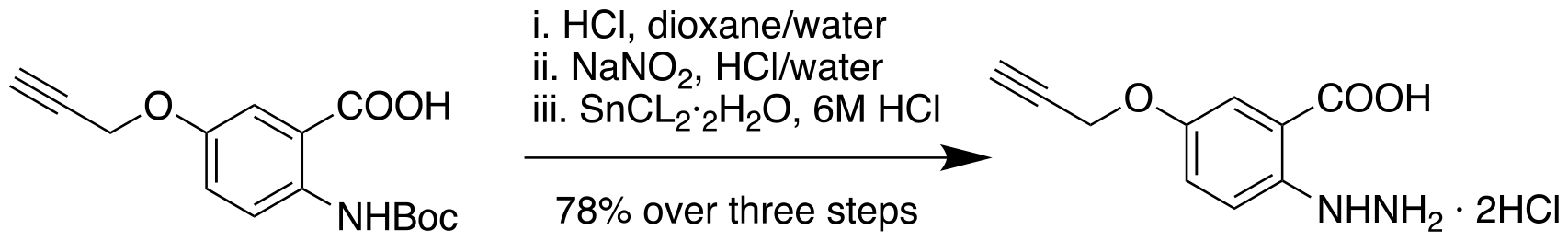
Synthesis of ProxyNA_3_

#### General synthetic procedures

Reagents were commercially available or prepared as stated below. All solvents were HPLC grade except for water (18.2 MΩ cm MilliQ water). Solvents were removed under reduced pressure in a rotary evaporator. Thin-layer chromatography (TLC) was carried out on Al backed silica gel plates with compounds visualised by anisaldehyde stain, 5% ninhydrin stain, and UV light. Melting points (mp) were obtained on EZ-Melt apparatus and are uncorrected. NMR spectra were recorded on a 300 MHz spectrometer for ^1^H NMR spectra δ values were recorded as follows: DMSO-D_6_ (2.50 ppm); for ^13^C (93.75 MHz) δ DMSO-D_6_ (39.50 ppm). Mass spectra (MS) were obtained using electron impact (EI).

#### ProxyNA_3_

Concentrated HCl (3 mL) was added to a stirred solution of 5-*O*-propargyl-*N*-Boc-anthranilic acid (291 mg, 1 mmol) in dioxane (2.5 mL). The mixture was stirred for 45 minutes at 80 ºC, was cooled to 0 ºC and an ice-cold solution of NaNO_2_ (76 mg, 1.1 mmol) in water (500 μL) was added. The stirring continued for 1 h at 0 ºC, an ice-cold solution of SnCl_2_ · 2H_2_O (451 mg, 2 mmol) in 6 M HCl (1 mL) was added, the cooling bath was removed, and the stirring continued for 1 h at room temperature. The mixture was set aside for 1 h at 3ºC, separated solid was filtered with suction, was washed with an ice-cold diluted HCl and was dried to afford 5-*O*-propargyl-*N*-aminoanthranilic acid dihydrochloride, orange solid, mp. 220-222 ºC, 217 mg, 78%. ^1^H NMR (DMSO-D_6_) δ 10.28 (br s, D_2_O exch., 3H); 8.82 (br s, D_2_O exch., 1H); 7.50 (d, *J* = 2.5 Hz, 1H); 7.30 (dd, *J* = 7.0, 2.5 Hz, 1H); 7.18 (d, *J* = 7.0 Hz, 1H); 4.81 (d, *J* = 2.0 Hz, 1H); 3.58 (t, *J* = 2.0 Hz, 1H). ^13^C NMR (DMSO-D_6_) δ 168.4, 150.9, 142.0, 122.0, 116.6, 116.5, 115.5, 79.1, 78.5, 56.0. HRMS (EI) *m/z*; found 206.0710 [M^+^] (calcd 206.0691 for C_10_H_10_N_2_O_3_). LRMS (EI) *m/z* (rel abundance) 206 [M^+^] (10), 131 (10). NMR spectra can be found in Supporting Figures 5 and 6.

### *In-vivo* Imaging

#### Dynamic Enhanced Contrast Magnetic Resonance Imaging (DCE-MRI)

The experimental design for DCE-MRI was adapted from a previous study^[29]^. The experiments were performed on a 3T MRI (MR Solutions). Mice were anesthetized with 1-2% isoflurane and then tail cannulation was performed for injection of the gadobutrol contrast agent. Single slice positioning was performed in the axial plane, between the bregma 3.56mm and -4.36mm. The following parameters were used: Slices: 7; Slice thickness: 1.0 mm; FOV: 40; Slice gap: 0.1; FOV ratio: 1.000. After setting the positioning parameters, and performing a Scout image, a series of serial T1-weighted images were obtained pre-contrast, during contrast, and post-contrast injection with gadobutrol (0.1 mmol kg-1; GadavistTM, Bayer). T1-weighted images were obtained using a Rapid Imaging with Refocused Echoes (RARE) pulse sequence with varying TIs (Pre-contrast: 280, 500, 900, 1200, 1500, 4650, 1200ms; During contrast: 1200ms x 10; Post-contrast: 280, 500, 900, 1200, 1500, 4650ms). The following imaging acquisition parameters were used: Echo spacing: 16; TE: 16; Echo train: 8; Views: 240.

Analysis of the scans was performed using the VivoQuant software. Four Regions of Interest (ROIs) were drawn using the 3D ROI Tool on 6 slices of each of the scans. Note, the first slice for each scan was not included in analysis due to the presence of the skull, which limited image resolution and clarity. The four ROIs included the brain, the ventricles, the meninges and the middle cerebral artery, which was referred to as blood. The mean voxel intensity in each of the ROIs across all six slices was generated for the scan immediately before gadobutrol injection (TI of 1200 ms) and then for the 10 consecutive scans taken at 3-minute intervals post gadobutrol injection (TI of 1200 ms). The data was then plotted for each of the ROI as the percentage change in voxel intensity, which was calculated by dividing the mean voxel intensity in each of the 10 consecutive scans taken post gadobutrol injection by the single scan taken immediately before gadobutrol injection. This procedure was performed for each of the timepoints: 5 days pre-impact (baseline), 6 hours post-impact, 30 hours post-impact, 54 hours post-impact. A generalized linear model (GLM), repeated measures was used to determine whether there were significant differences in the percentage change in voxel intensities in each of the ROIs before and after impact.

#### CEST-MRI

All CEST-MRI acquisitions were performed on a 3T pre-clinical MRI (MR Solutions, Ltd.). A single slice Single Shot Rapid Imaging with Refocused Echoes (RARE) pulse sequence was implemented with the following parameters: Average=1, Matrix size=96x96, TE=7 ms, Echo Spacing=7 ms, TR=6000 ms, CEST Length=5000 ms.. Prior to imaging, tail veins were cannulated with an ∼1m long cannula (∼100 μL dead volume), permitting *intravenous* injection in the MRI. Mice were warmed and respiration rate was monitored through the duration of the imaging experiment. Heads were centered in the FOV using standard localizer sequences. In order to correct for B_o_ inhomogeneity, a Water Saturation Shift Referencing (WASSR) sequence was implemented prior to any Z-spectra acquisitions. For WASSR, Z-spectra were acquired from +200 to -200 Hz at 25 Hz steps, and a pre-saturation pulse B_1_ amplitude B_1_ = 0.65 μT . The total acquisition time for WASSR was 1 min 42 seconds. CEST MRI prior to contrast agent administration (single slice, slice thickness = 2.0 mm FOV = 35 x 35 mm, CEST frequency range from +1500 to -1500 Hz centered on the water resonance, sampled in steps of 100 Hz, a pre-saturation pulse B_1_ amplitude B_1_ = 2.61 μT, and acquisition time 3 minutes 6 seconds). After the completion of the initial Z-spectrum, each mouse received a bolus injection of 100 μL of 40 mM of PHBA dissolved in saline, and the line was flushed with 100 μL saline. No adverse events were noted in any of the mice receiving PHBA injections. Post-contrast Z-spectra were acquired sequentially for 45 min post-injection. An anatomical reference image was acquired using a T_2_-weighted RARE sequence (Average=2, Matrix size=256x256, TE=68 ms, Echo Spacing=17 ms, TR=4800 ms) with the same FOV as the Z-spectra acquisitions.

Images were post-processed and analyzed using the Matlab routine provided by Guanshu Liu et al., available for download here: http://godzilla.kennedykrieger.org/CEST/ ^[30]^. Code modifications were necessary to read the vendor specific imaging files and to reorder the image data. Image maps were generated to plot %MTRasym at Dw = 6.5 ppm over the entire slice, and a final map of the ratio of 45 min post-PHBA injection divided by pre-PHBA injection images was plotted using Image J. This map was colour coded using a custom lookup table and overlaid onto the corresponding anatomical image. Brain regions of interest (ROI) of the ratio map were quantified by histogram analysis using ImageJ.

### Histology

Mice were anesthetized with isoflurane and perfused with 4% PFA. The brain was delicately removed from the skull and placed in a 50mL tube containing 4% PFA, at 4°C for 24 hours. The brains were then cryoprotected in sequential 15% and 30% sucrose solutions at 4°C. They were then embedded in OCT (Tissue-Trek) over dry ice and moved to -80°C for storage.

#### Tissue sectioning and free-floating immunofluorescence staining

Serial 40µm sections were taken using a cryostat (Leica CM1850) starting at the initial appearance of the lateral ventricles and placed into a 24 well plate containing PBS with 0.01% sodium azide solution and stored at 4°C. 12 sections were taken per brain.

12 brains were excised from mice from each group (naïve, 2 days post-injury, 7 days post-injury) for immunofluorescence staining. One brain was excluded from the 7-day group due to damage during excision. Therefore, 35 mice were used for histological assessment.

Brain sections were removed from sodium azide-containing solution and washed in PBS 3x 5min on a rocker. They were then blocked with a solution containing 5% goat serum, 0.1% Triton X-100 and 0.1% Tween-20 for 1 hour at room temperature. The sections were then incubated with primary antibodies (α-GFAP or α-Iba1) diluted in blocking solution overnight at 4°C on a rocker. After washing with PBS 3x 5min, secondary antibodies were added in blocking buffer for 1 hour at room temperature on a rocker. Secondary antibody solution was removed, and the sections were washed again and then transferred into a petri dish filled with 1X PBS and mounted onto a charged microscope slide using a fine paintbrush. A coverslip was placed onto the slide using Fluoromount-G mounting medium with DAPI (Invitrogen), and the slide was stored in the dark until imaging.

Imaging was performed on a Zeiss axioscan microscope slide scanner. Images were tiled, z-stacks were taken to ensure total depth was imaged, and at 10X resolution. Post-processing in Zen Blue included tile stitching, shading correction, background subtraction, and maximum intensity projection. For quantification, a 1500x1300pixel rectangular region of interest was drawn on approximately the same region of hippocampus of each image and quantified in Fiji (total GFAP intensity divided by total Hoechst intensity).

#### Western Blotting

Mice were euthanized via CO_2_ asphyxiation. Brains and livers were removed and immediately snap-frozen in liquid nitrogen and stored at -80°C until use. Tissues were then homogenized in 1X RIPA buffer with protease inhibitor tablet (Roche) in a bead beater for 5 minutes at 4°C. The protein in the supernatant was then quantified using Pierce BCA protein quantification kit with BSA in lysis buffer for standard curve. 30µg of protein was added per well and run on a 4-20% pre-cast gel (Biorad). Membranes were probed with α-ALDH2 (Proteintech 15310-1-AP) and α-Vinculin (Abcam ab130007).

## Results

### Imaging aldehydes *in vitro* in a 3D corticomimetic scaffold after concussive impact

A three-dimensional collagen-based scaffold was prepared to mimic the biomechanical (i.e., stress/strain) properties of mouse cortical brain tissue. The corticomimetic scaffold consisted of two parts (Fig. 1A, top): a stiffer outer ring comprised of glutaraldehyde-crosslinked collagen necessary to mimic mouse cortical brain mechanics and provide overall structural support to the construct; and a gel-like inner plug comprised only of collagen, which optimally promoted the growth and survival of stem cell-derived mammalian cortical neurons (Fig. S1). In order to mimic mouse cortical brain tissue, the Young’s Modulus of a fresh punch of mouse cortical tissue was determined using a universal testing machine (UTM) (Fig. 1B). We determined that a corticomimetic scaffold comprised of an outer ring of 8.1 mg/mL collagen + 0.75 wt% glutaraldehyde, and inner plug of 2 mg/mL collagen best mimics the biomechanical properties of mouse cortical brain (Fig. 1B). In maintaining the formulation of the inner plug at 2 mg/mL collagen with no cross-linker, the growth of primary neural stem cell-derived mouse cortical neurons (Fig. 1C and D) and induced pluripotent stem cell (iPSC)-derived human cortical neurons (Fig. S2) was promoted. Using NeuO as a fluorescent neuronal marker ^[31]^, confocal microscopy revealed both mouse (Fig. 1C) and human (Fig. S2) neurons had migrated throughout the scaffold, resulting in neurite outgrowth. The application of live/dead staining confirmed the cells were alive, with very minimal death seen after seven days post-seeding (Fig. 1D and Fig. S2).

**Fig. 1.**
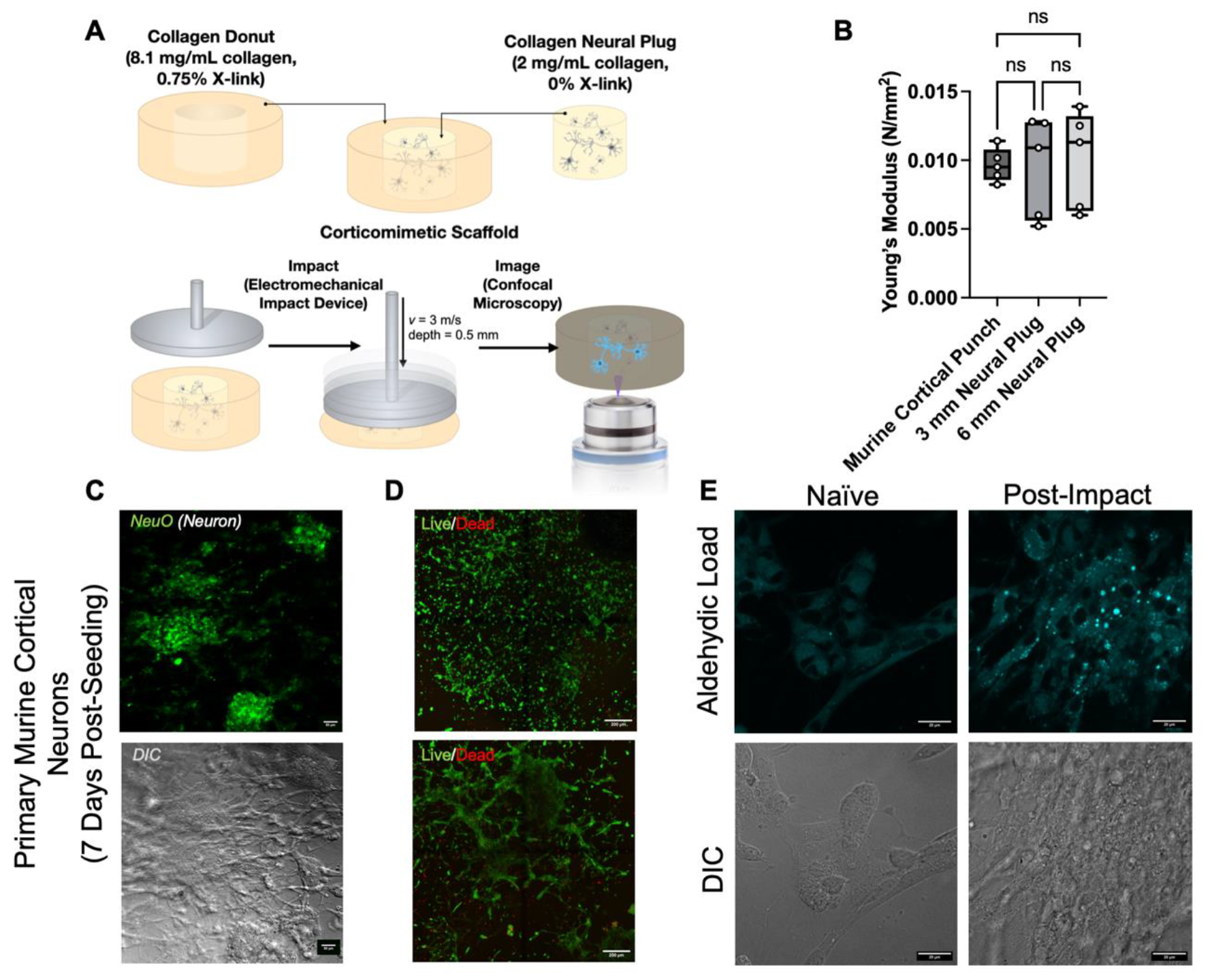
Aldehydes are produced following concussion-like impact in primary neurons growing in a corticomimetic scaffold. (**A**) Formulation of corticomimetic scaffold with crosslinked collagen exterior, and collagen interior containing primary mouse neurons. The scaffold was impacted with a concussion-like force followed by confocal microscopy with an aldehyde-binding fluorophore 5MeONA_3_ (**B**) Young’s Modulus of the corticomimetic scaffold with two different sized interior plugs compared to a murine cortical punch. Data are shown in a box plot with individual values, n=5 per group, p<0.05. (**C**) Confocal microscope images showing NeuO fluorescent staining (top) of primary mouse cortical neurons seven days after seeding into corticomimetic scaffold to confirm differentiation of neurons. Differential interference contrast (DIC; bottom) shows elongation of cells within 3D scaffold. Scale bar = 50µM (**D**) Evaluation of cell death at three- and seven-days post-seeding of primary mouse cortical neurons into corticomimetic scaffold. Scale bar = 200µM. (**E**) Representative confocal images of primary neurons pre- (left) and post- (right) concussion-like impact after incubation with 5MeONA_3_ for fluorescent staining of aldehydes (top) and DIC of cells showing morphological changes pre- and post-impact. Scale bar = 20µM.

With the ability to support primary neuron culture in a scaffold recapitulating mouse cortical brain rheological properties, the corticomimetic scaffold was then applied for biomarker testing post-concussion (Fig. 1A). A custom impactor tip affixed to an electromechanical impactor (Leica Inc.) produced a force to compress but not tear the scaffold, and to mimic a previously-reported concussion-like strain of 26% thickness ^[32]^. A cell-penetrable, aldehyde-conditional fluorogenic probe (5MeONA_3_) ^[26]^ afforded the mapping of total aldehydic load in the corticomimetic scaffolds by confocal microscopy before and 10 minutes post-impact (Fig. 1E). Every post-impact scaffold exhibited an increase in aldehyde signal over the baseline, pre-impact levels present as a result of normal metabolism. Interestingly, the aldehydes presented as round, punctate foci within the cytosol (Fig. 1E). In primary neurons cultured in a corticomimetic scaffold, total aldehydic load increases after a concussive-like impact, supporting the role for aldehydes as a biomarker for cell injury following mTBI.

### Development of an awake closed head impact mouse model of mTBI

The use of an awake, closed head impact (ACHI) mouse model of concussion is important to provide behavioural and neuroimaging outcomes most relevant to human injury. Many rodent models of TBI have been developed ^[33]^, however surgical manipulation, head fixation, and the use of neuroprotective anesthesia make these models less translatable to humans. Linear and rotational acceleration of the head are a fundamental characteristic of mTBI ^[34]^, which is lost with fixed-head or surgical models. The use of anesthesia provides neuroprotection ^[35]^, which alters the biochemical and behavioural outcomes of the impact. Furthermore, an awake model provides the opportunity to perform behavioural testing immediately after injury, which is a crucial timepoint as many symptoms of concussion in humans are commonly identified immediately after impact ^[36]^. By adapting two previously described procedures ^[37]^, a new ACHI mouse model of mTBI is developed, eliminating limitations of surgical, fixed head, and anesthetized models.

Impact velocity is a crucial variable for a reproducible mTBI. Velocities of 2.35 to 6m/s have been previously described in the literature (Table S1). A suitable ACHI model would elicit some behavioural phenotype, but cause no blood-brain barrier (BBB) leakage, no skull fracture or intracranial hemorrhaging, and no severe reaction including seizure, paralysis, or death ^[38]^. The ACHI model was initially developed on 8–10-week-old healthy, wild type (WT) Balb/c mice (Fig. S3A). A single impact with velocities of 2.35, 5.0, or 6.0 m/s were evaluated (n=5 mice/group). Mice receiving impacts at 6.0m/s showed paralysis of limbs, inability to regain righting reflex, and intermittent apnea. Post-mortem dissection revealed skull fracture and intracerebral hemorrhaging of the right cortex, indicating 6.0 m/s resulted in TBI, not mTBI. No such damage was observed during post-mortem gross evaluation of the brains of mice receiving 2.35 or 5.0 m/s impacts.

As behavioural deficits are a hallmark of rodent concussion ^[33a]^, grid walk (Fig. S3B,C) and neurological severity score (NSS; Fig. S3D-H) tests were conducted before impact, immediately after regaining righting reflex following impact, and at 1-, 4-, and 24-hours post-injury (Fig. 2 A-C). NSS consists of a series of 5 objective pass or fail tests used to assess balance, cognition, reflexes, and exploratory behavior. A higher score indicates a greater deficit, with a score of 5 being the most severe. Sham mice (restrained with no impact) had a baseline average score of 0.2±0.4, which did not significantly change at any evaluation timepoint . The NSS of mice receiving an impact of 5m/s significantly increased to 3.8±0.4 (p<0.05) immediately after impact and returned to baseline for all subsequent time points (average 0.8 to 1.2; p>0.05) (Fig. 2A). There was no change in NSS for mice receiving 2.35 m/s impact. The grid walk test assesses acute deficits in motor coordination and overall activity level ^[39]^. The percentage of foot faults compared to number of total forelimb steps and percent time active was counted from a 3-minute video (Movies S1,S2). For mice receiving a 5 m/s impact, forelimb foot-faults increased significantly over baseline (18.8±16.4% *vs*. 6.0±0.19%, p<0.05) (Fig. 2B), and activity time decreased significantly (40.6±17.6% *vs*. 85.8±25.4%, p<0.05) immediately after impact (Fig. 2C).

**Fig 2.**
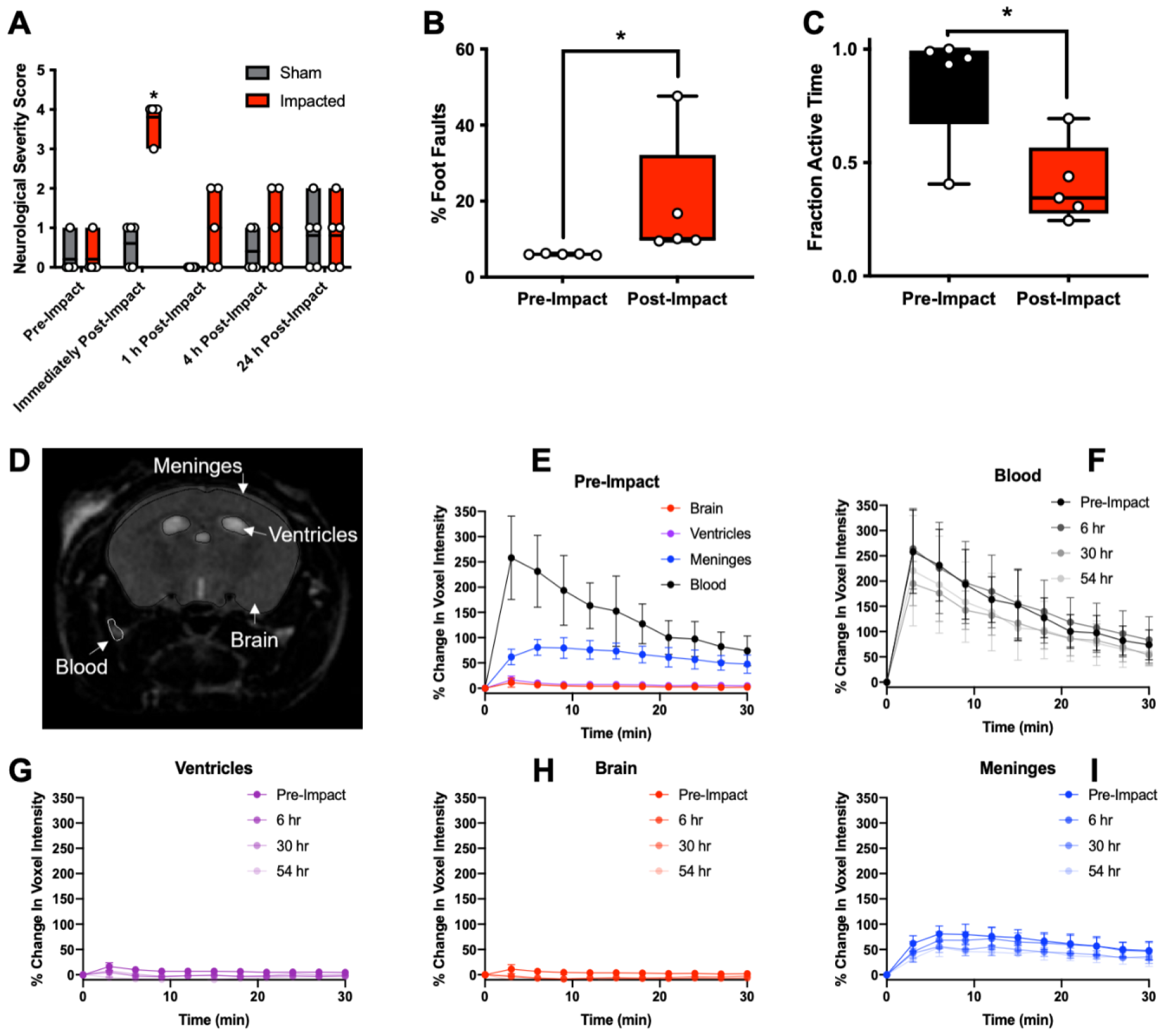
Mice exhibit changes in behaviour but not blood-brain barrier permeability after mTBI. (**A**) Mice underwent behavioural testing for neurological severity score (NSS) comprised of 5 pass-or-fail tests including beam walk, hindlimb reflex, startle reflex, escape paradigm, and balance test. A point was awarded for every test failed, with a score of 5 representing the more severe behavioural deficit. Individual data points (n=5 per group) and box plots are shown. Sham mice were restrained, but not impacted. * p<0.05. (**B-C**) Results from 3-minute grid walk test for pre- and post-impact mice (n=5 per group). (B) Percentage of foot faults of total forelimb steps before and after ACHI. (C) Time active including walking, grooming, and exploratory behaviour during the three-minute grid walk challenge is shown. Statistical analysis was performed using a One-Tailed Wilcoxon Test, * denotes p< 0.05. (**D**) Regions of interest (brain, ventricles, meninges, and blood) were drawn to evaluate contrast enhancement following intravenous gadolinium-based contrast agent pre- and post-impact. (**E-I**) Dynamic contrast enhanced MRI was performed pre-and post-impact at various timepoints. Percent change in voxel intensity after gadobutrol injection in each of the ROIs from images acquired (E) five days pre-impact and (F-I) at timepoints of 6 hours-, 30 hours-, and 54 hours post-impact. Data are shown as means +/- standard deviation. Each data point represents the analysis of ROIs on 6 slices per brain MRI. P>0.05 using GLM repeated measures, n=5 mice.

Following the observation of behavioural sequelae to ACHI at 5.0 m/s, BBB integrity was assessed with DCE-MRI ^[40]^. Four regions of interest were analyzed for contrast enhancement pre- and post-impact: ventricles, middle cerebral artery as a proxy for blood, the meninges, and the remaining brain structures (Fig. 2D) ^[41]^. Pre-impact, blood contrast increased (246%) (Fig. 2E), with no significant change in contrast enhancement in the brain tissue or ventricles (p>0.05). There was a significant increase in contrast in the meninges by 6 minutes after gadobutrol injection, however the lack of extravasation, retention in the subarachnoid space and steady clearance over time signifies an intact BBB ^[40b]^. Post-ACHI, there were no significant changes in contrast enhancement in blood (Fig. 2F), ventricles (Fig. 2G), brain (Fig. 2H), and meninges (Fig. 2I) relative to pre-impact, suggesting ACHI does not cause disruption of BBB integrity, supporting the mild severity of our model recapitulating concussion. Overall, we have shown that a single-impact ACHI mouse model with impact velocity of 5.0m/s, depth of impact of 7mm, and dwell time of 0.3s results in transient behavioural deficits without overt signs of skull damage, without intracranial hemorrhage, and without compromising BBB integrity.

### CEST-MRI mapping of total aldehydic load *in vivo* following mTBI

We have previously reported *N*-aminoanthranilic acids as aldehyde-conditional CEST-MRI contrast agents, where MRI contrast is produced only upon reaction of the probe with aldehydes to form a hydrazone (Fig. 3A) ^[42]^. The reactive *ortho* acid hydrazine moiety has been shown to effectively bind aldehydes *in vivo* ^[27, 43]^, and here we introduce ProxyNA_3_ to map aldehydic load after mTBI. Two hypotheses were evaluated in the current study: (1) brain aldehydic load increases in aged relative to younger mice following concussion; (2) ALDH2 is involved in concussion-related pathogenic aldehyde metabolism. In order to evaluate these hypotheses, WT C57Bl/6, and previously reported C57Bl/6 mice harboring heterozygous or homozygous knockouts of ALDH2 were employed ^[20]^. ALDH2 is normally expressed in the liver but not the brain ^[44]^ (Fig. 3B). Within these three groups, mice were evaluated when either young (<20 weeks) or of advanced age (>65 weeks). Four male mice were induced to ACHI from each cohort (total of n=24 mice) and underwent behavioural testing at two timepoints (pre- and post-impact), and *in vivo* MRI at three timepoints (pre-injury, two days-, and seven days-post injury) (Fig. 3C). An additional 6 mice from each group were used for histological evaluation at each of pre-injury, two days-, and seven days-post injury (n=36) (Fig. 3C).

**Fig 3.**
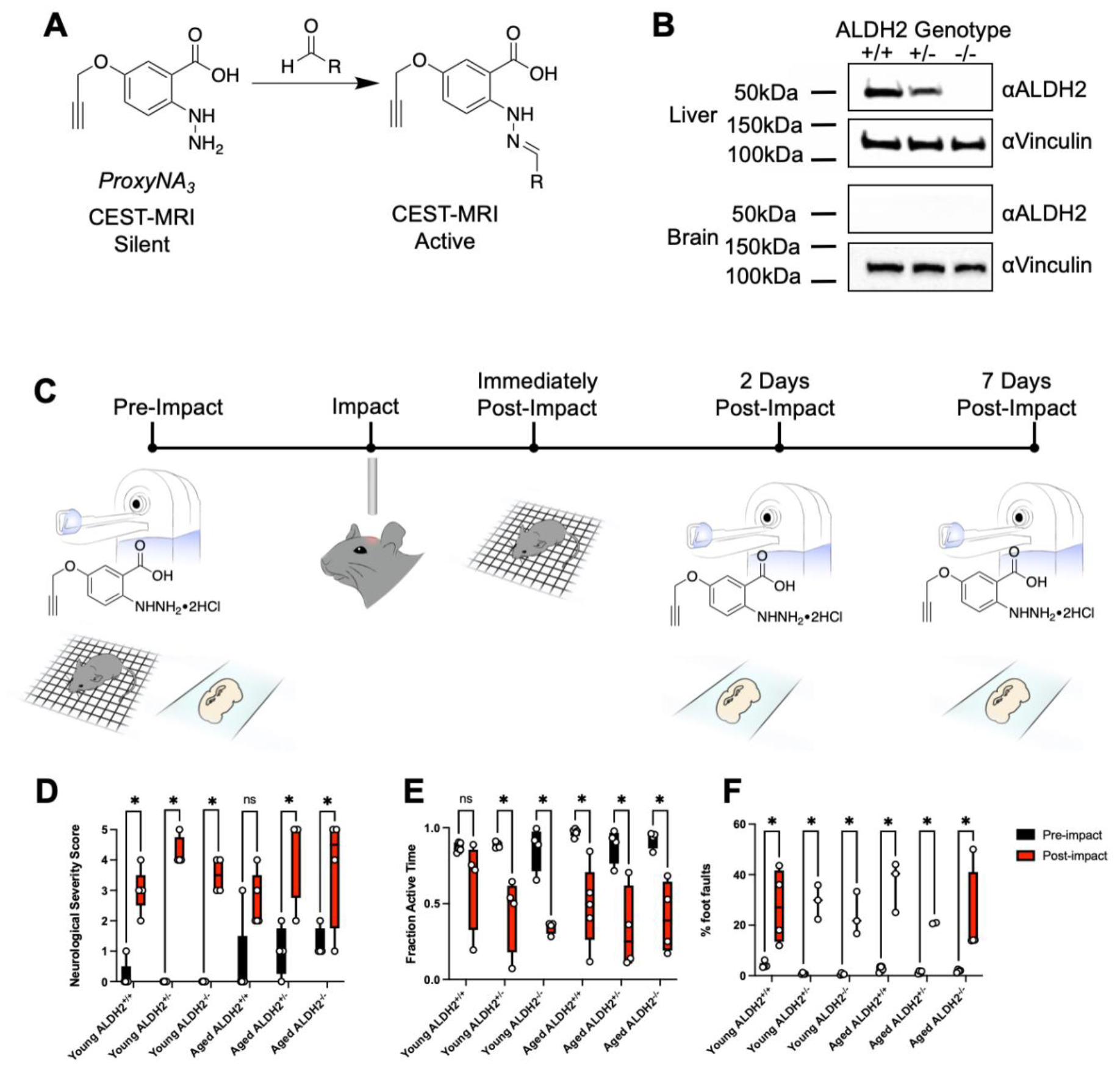
Experimental procedure for evaluation of mTBI in ALDH2^-/-^ mouse model using behavioural tests, histology, and ProxyNA_3_ CEST-MRI. (**A**) Chemical structure of ProxyNA_3_ in its unbound, silent state (left), and its CEST-MRI active state, upon reaction with aldehyde (right). (**B**) Western blots showing protein levels of ALDH2 in the liver (top) and brain (bottom) tissues from transgenic mice with ALDH2 ^+/+, +/-^, and ^-/-^ genotypes. Vinculin was used as a loading control protein. (**C**) Experimental design for all testing done pre- and post-impact for *in vivo* mouse model of mTBI. CEST-MRI with ProxyNA_3_ was performed pre- and at two- and seven days post-impact. Behavioural tests, denoted by grid-walk symbol, was performed pre- and immediately post-impact. A separate cohort of mice were used for immunohistological examination at the same timepoints as MRI. (**D**) Results of neurological severity score (NSS) comprised of 5 pass-or-fail tests pre-impact (black) and post-impact (red). A point was awarded for every test failed, with a score of 5 representing the most severe behavioural deficit. (**E-F**) Immediately after NSS test, mice were placed on a raised grid with video recording for 3 minutes. A researcher blindly scored (E) the fraction of time spent active and (F) the total number of foot faults divided by total number of forelimb steps. Groups of mice were chosen by age (young <16 weeks; aged >65 weeks) and by genotype. Individual data points (n=4 per group) and box plots are shown. * p<0.05, 2-way ANOVA followed by Tukey-corrected multiple comparisons test.

All mice had an increased NSS immediately after impact compared to baseline (Fig. 3D). The increase was statistically significant in all cohorts except for the aged WT mice (p=0.063). All mice had a decreased fraction of time spent active during the grid walk (Fig. 3E), with only the young WT cohort failing to reach statistical significance(p=0.749). Every group had a significant increase in number of foot faults on the grid walk following injury compared to baseline (Fig. 3F). The results of the behavioural testing confirmed transient sensorimotor deficits indicative of mTBI across all groups of mice tested.

CEST-MRI was performed before and after injection of ProxyNA_3_ to map total aldehydic load in a single axial slice of the brain. While brain aldehyde levels were low before impact (naïve), normalized %MTR_asym_ increased significantly over the whole brain slice and vasculature in all mice at two-days post-impact (Figs. 4, S4). However, by day seven post-impact, aldehyde signal decreased to baseline levels, suggesting that aldehydes are transiently present in the brain following mTBI in mice. Young WT mice have less aldehyde signal compared to all other groups (Fig. 5A). Aged WT mice have less aldehydes compared to all other aged mice, with both cohorts of aged ALDH2 deficient mice having the greatest aldehyde signal two days post-injury. When grouped by age-only, young mice have significantly less aldehyde-derived imaging signal compared to aged mice two days post-injury (Fig. 5B). ALDH2 deficient mice (both ALDH2 ^+/-^ and ^-/-^) had significantly more aldehyde signal compared to WT mice (Fig. 5C). These data suggest that more toxic endogenous aldehydes are produced following mTBI in aged individuals, and those with ALDH2 deficiency.

**Fig 4.**
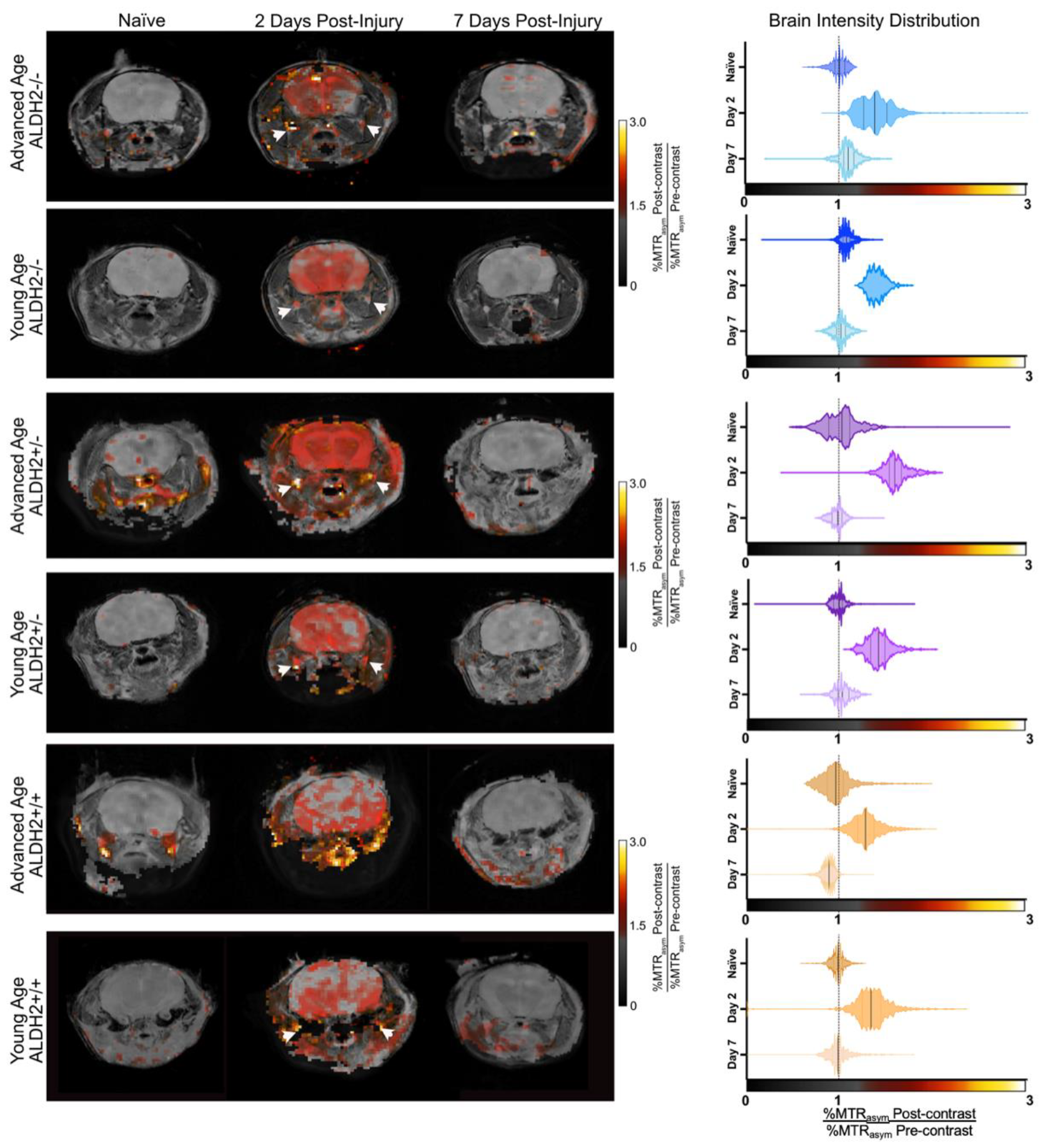
Mapping of brain aldehydes by ProxyNA_3_ in a mouse model of mTBI. CEST-MRI (average %MTR_asym_ normalized to pre-contrast signal) after ProxyNA_3_ injection mapped onto axial T_2_-weighted image of mice of identified age group and ALDH2 deficiency (left). A representative image of a mouse from each cohort is shown longitudinally across timepoints (pre-impact, two days post-impact, and 7 days post-impact). Arrowheads indicate vascular signal derived from ProxyNA_3_-aldehyde adducts. Violin plots showing voxel-wise intensity distribution for the corresponding mouse at each timepoint. Solid line= median, dashed lines = 25^th^ and 75^th^ %ile.

**Fig 5.**
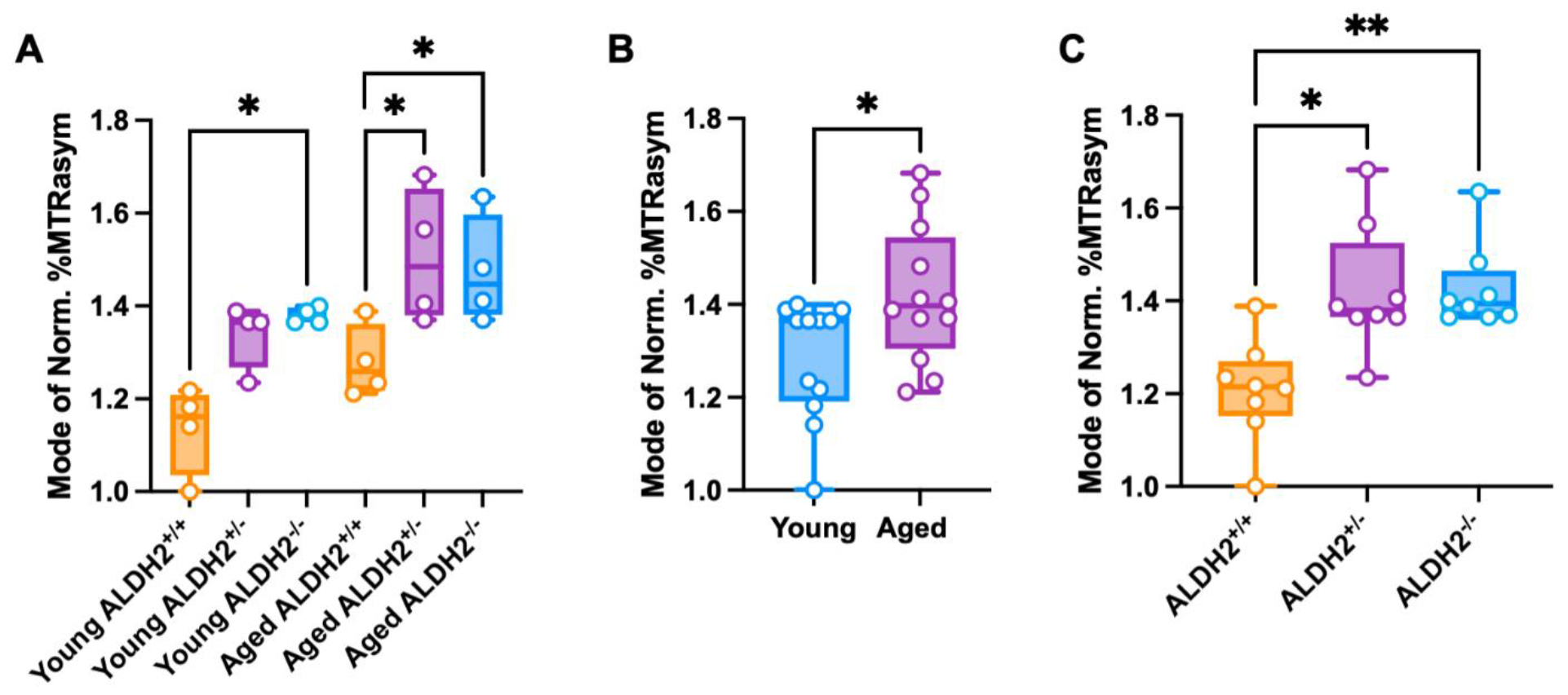
Aldehyde production post-concussion is affected by age and ALDH2 genotype. Box plots of voxel-wise intensity distribution of aldehyde signal following ProxyNA_3_ CEST-MRI at day two post-mTBI. Mice are grouped by age and genotype. Data are presented as modes of normalized %MTR_asym_. (**A**) Comparison of aldehyde signal two days post-impact between all groups of mice, n=4 per group. (**B**) Comparison of young (<16 weeks) and aged (>65 weeks) mice, n=12 per group. (**C**) Comparison of aldehyde signal between groups of either wild type (ALDH2^+/+^) or transgenic mice with lower ALDH2 expression (ALDH2^+/-^), or complete knockouts (ALDH2^-/-^). *<0.05, (A, C) Brown-Forsythe ANOVA and Dunnett’s T3 multiple comparison, (B) two-tailed Mann-Witney U-test.

### Aldehydes precede astrogliosis in the pathogenesis of mTBI

Astrocyte activation, an active proponent in the propagation and regulation of neuroinflammation ^[45]^, was quantified by immunohistological staining for glial fibrillary acidic protein (GFAP) in brains of mice pre- and at two timepoints post-injury (Fig. 3C). Hippocampal astrogliosis appeared seven days post-injury for both young mice (Fig. 6A) and those of advanced age (Fig. 6B) independent of ALDH2 genotype. A region of hippocampus from each image was quantified by measuring GFAP density, the mean integrated density of GFAP signal divided by the mean integrated density of DAPI signal, which estimates the astrocyte activation normalized to number of cells (Fig. 6C.). There was no significant difference in GFAP density by age and by genotype. The only variable influencing the measure being time relative to ACHI, with GFAP density increasing from 1.9±0.26 to 2.7 ± 0.78 (p<0.05) at 7-days post-impact. These results suggest an activation of a hippocampal inflammatory response seven days after injury, proceeding the elevation of aldehydic load (Fig. 6D).

**Fig 6.**
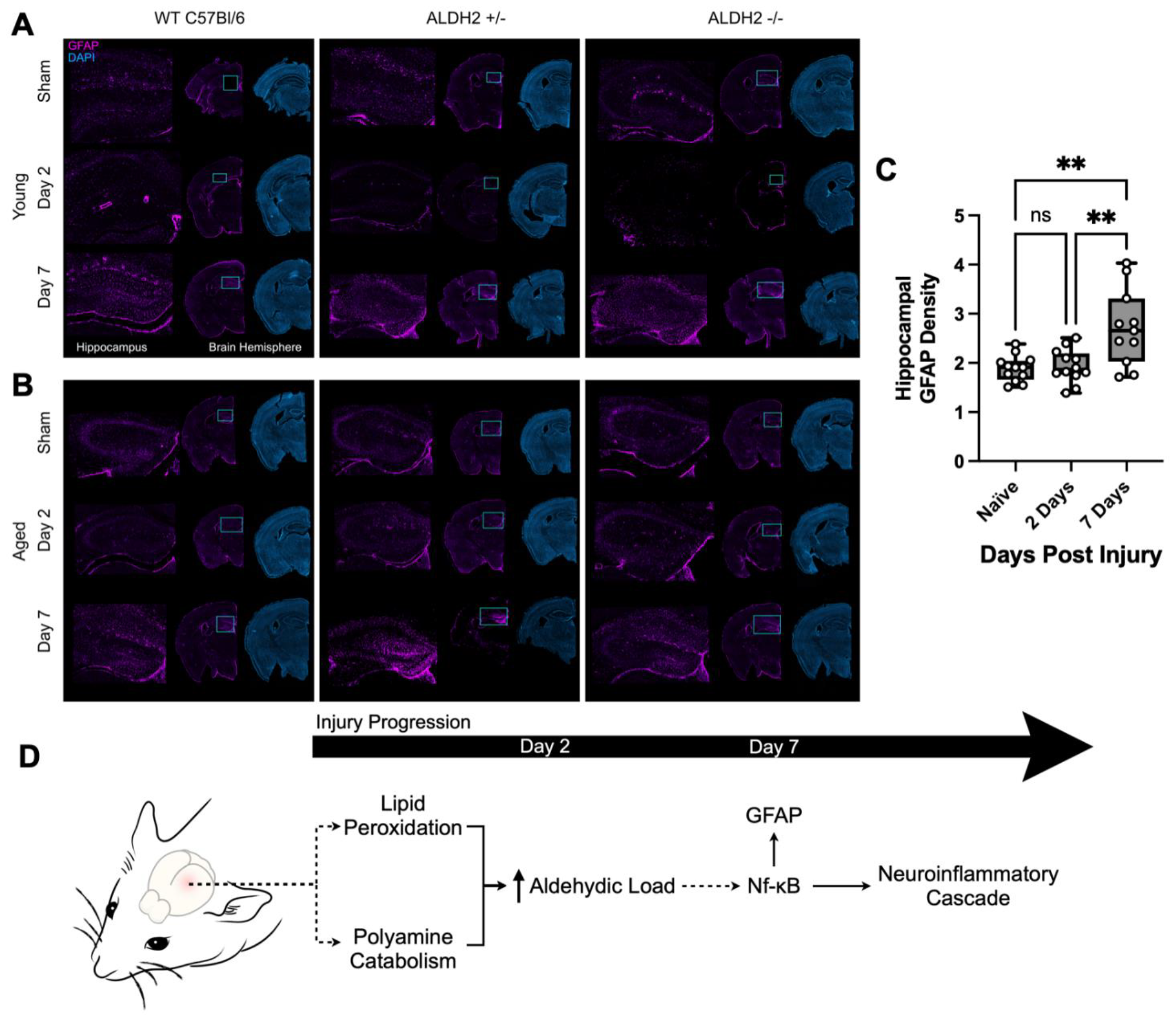
Astroglial activation increases 7 days post-impact, following increase in aldehydic load during the neuroinflammatory cascade driving mTBI. The brains of mice were excised at day two- and day seven-post-impact, or without impact (sham). Brain sections were fluorescently probed with anti-GFAP antibody to visualize astrocyte activation and DAPI. A representative image is shown for a mouse in each cohort based on genotype, age, and timepoint. (**A**) Young mice (<16 weeks) and (**B**) old mice (>65 weeks) all show an increase in GFAP signal in the hippocampus at seven days post-injury, but not at two days post-injury compared to sham mice. (**C**) Quantification of GFAP density in a region of the hippocampus from each mouse with no impact, or two- and seven-days post-impact (n=12 per group; **p<0.01, ns, non-significant, one-way ANOVA with *post-hoc* Tukey’s test. (**D**) Proposed causative role of aldehydes, produced from oxidative events, in the pathogenesis of mTBI: initial creation of aldehyde-rich tissue, followed by astrogliosis (Nf-κB-driven GFAP expression), and progression of the neuroinflammatory cascade.

## Discussion

Our corticomimetic scaffold was designed to mimic the mechanical properties of cortical brain tissue upon compression (Fig. 1B), and support the growth of primary neurons using existing mammalian cell culture infrastructure and commercially-sourced collagen. The collagen-based 3D scaffold offers some practical advantages over bioprinted or silk-based scaffolds ^[46]^ by reducing infrastructure-need, process complexity, and time-to-finished product ^[46b, 47]^. This higher-throughput, cost-effective method may improve *in vitro* testing of CNS-injury models, and, through co-culture of neurons with astrocytes and microglia, provide a more complete examination of the neurometabolic cascade following injury. In the current study, this scaffold allowed for imaging of aldehydic load in primary mouse cortical neurons before and after concussion-like impact. Naïve, unimpacted cells have some background level of aldehydes due to normal metabolic processes, consistent with our previous findings ^[26]^. Following impact, the cells have a marked increase in aldehyde-rich foci within the cytoplasm of cells (Fig. 1E), likely due to both lipid peroxidation and polyamine catabolism ^[10e, 10f, 26]^. Future work will aim to determine the nature of these aldehyde-dense foci.

In recent years, one focus of TBI diagnostic research has been the discovery of blood-based biomarkers, including several astroglial and neuronal biomarkers S100B, GFAP and UCH-L1 ^[48]^. Such blood tests provide no spatial dimension to the diagnosis (i.e., implicated brain regions) and only indicate that an injury has occurred. Molecular imaging of aldehydes fills this gap, providing information about severity, and location of injury, which will not only diagnose with high sensitivity, but will aid in the prognosis and personalized treatment planning following mTBI. The ALERT-TBI study identified a prespecified cut-off value for serum GFAP and UCH-L1 to predict intracranial injury on head CT ^[49]^, which contributed to the FDA approval of the first blood test for avoidance of unnecessary CT in adults following head trauma in 2018. Studies on serum and plasma GFAP to discriminate mTBI with and without CT abnormalities are emerging ^[50]^, with GFAP continuing to outperform other biomarkers ^[51]^. Although blood GFAP level shows promise, there are some limitations regarding its utility for concussion diagnosis. GFAP levels in the blood may be sensitive to intracranial pathologies that are not seen on CT, and there is continuing debate on how the mechanisms by which GFAP and its breakdown products exit the cell and enter the bloodstream. Importantly, GFAP expression increases with age in the healthy population ^[52]^, reducing the specificity of a blood-based test in older individuals ^[53]^. In general, fluid biomarkers have limitations, including short half-life in biofluid, sources for generation outside of the CNS, delayed appearance in the circulation, and dependence on age, sex, and co-morbidities ^[54]^.

ALDH2 deficiency affects 560 million people worldwide, making it one of the most prevalent hereditary disorders ^[55]^. The results from this study indicate a potential increased risk for more severe outcomes for the population with ALDH2 deficiencies and those of advanced age. Mice with ALDH2 deficiency had a larger increase in aldehydes two days post-mTBI compared to WT mice (Fig 5. A, C). Furthermore, aged mice had more brain aldehydes post-injury compared to young mice (Fig. 5 B). Previous studies have shown that aged mice have decreased ALDH2 activity and increased HNE (4-hydroxynonenal) protein adducts compared to young mice ^[56]^, which exacerbates the increase in mitochondrial ROS production and accumulation of aldehydes that comes with age ^[57]^. Together, these findings support our imaging results to suggest that the ability to clear these aldehydes is decreased with reduction in ALDH2 activity due to a genetic mutation (ALDH2 deficiency) and/or advanced age. If aldehydes are propagators of disease, the inability to quickly clear aldehydes produced in the brain following mTBI may exacerbate post-impact pathophysiology and/or delay recovery. Now that aldehydes have been determined to be an early biomarker for mTBI, further testing should be done to elucidate the pathogenic role of aldehydes in the neurometabolic cascade and if there is an increased risk of more severe symptoms, including PCS, following mTBI in the aged and ALDH2-deficient populations.

It is known that oxidative stress leads to profound activation of NF-κB, a family of transcription factors that play important roles in stress responses, inflammation, and apoptosis ^[58]^. Various oxidants, cytokines, chemokines, and growth factors induce molecular signals that lead to activation of NF-κB ^[59]^, a central step in pro-inflammatory reactive astrocytosis ^[60]^, reactive gliosis, and cellular atrophy ^[61]^. Several studies have shown a link between NF-κB and GFAP upregulation following TBI in cells and *in vivo*, with NF-κB-induced astrocyte activation driving edema ^[62]^. Interestingly, inhibition of ALDH2 activity results in enhanced nuclear translocation, and therefore activation, of NF-κB ^[63]^. Inhibition of NF-κB has also been shown to reduce astrogliosis in mice ^[64]^. While it is not clear exactly how oxidative stress activates various protein kinases upstream to NF-κB, evidence implicates stress-derived aldehydes in the activation of the signaling cascade that eventually activates NF-κB ^[65]^. We propose that the link between initial mTBI-induced injury and NF-κB signaling is mediated by aldehydes as secondary messengers. Lipid peroxidation-derived aldehydes are known to act as toxic secondary messengers to propagate redox signals leading to cellular and tissue injury ^[66]^. Our data directly support the hypothesis (Fig. 4 and 6) that mTBI-induced aldehyde production precedes GFAP overexpression, the result of NF-κB activation, which contributes to reactive gliosis and inflammation in the brain to propagate trauma-induced injury.

A limitation of the current study is that the earliest timepoint for *in vivo* aldehyde mapping was two days post-injury. CEST-MRI will be repeated at shorter post-injury intervals to assess how early acute aldehyde production occurs. Our study did not explore sex differences in aged and ALDH2^-/-^ mice. Some literature suggests ALDH2 polymorphisms may affect women more than men ^[67]^, although the effect of sex is likely different in mice compared to humans ^[57]^. Moreover, the CEST-MRI technique used for *in vivo* imaging was limited to a single slice, preventing whole-brain analysis. Efforts are ongoing to translate the ProxyNA_3_ probe chemistry described here to a radiotracer for positron emission tomography (PET). Not only would a radiotracer similar to ProxyNA_3_ described herein allow for quantitative whole-body mapping of aldehydic load following mTBI, but its successful implementation could be more rapidly translated to human use ^[68]^.

The current study represents the first steppingstone in the application of aldehydic load as a biomarker for the objective diagnostic imaging of mTBI. Here, we propose a novel 3D *in vitro* model of concussion, which supported the use of aldehydes as a biomarker for mTBI. A new mouse model of ACHI was developed resulting in behavioural and histological sequalae recapitulating mTBI. Using this model, ProxyNA_3_ CEST-MRI was used to map the increase in aldehydic load in the brain following concussion. Future iterations of this imaging tool may be used in a clinical setting not only telling that mTBI occurred, but also mapping the severity and location of damage, and leading to improved treatment outcomes *via* personalized concussion medicine.

## Acknowledgements

The authors would like to thank Dr. Emilio Alarcon for his advice on preparation of 3D scaffolds, and Vivian Franklin for her assistance in mouse colony breeding and maintenance.

## Funding

A.J.S. acknowledges funding support from Canada Research Chairs (950-230754), CIHR (PJT-376892), NSERC Discovery Grant (RGPIN-2021-03387), Government of Ontario Early Researcher Award (ER18-14-102), and the Concussion Injury Group of the uO Brain and Mind Research Institute.

## Author contributions

A.K., R.D., and A.S. developed *in vivo* mouse model of mTBI. A.K., C.W., A.S. and N.D.C. contributed to corticomimetic scaffold development. J.W. and J.L. performed primary cortical precursor cell dissections. M.S. and N.C. performed all chemistry for probe preparation. A.K., R.D., C.D., and E.S. conducted UTM experiments. A.K. performed histology, microscopy, and molecular biology procedures. A.K. and A.J.S. conducted MR imaging, performed statistical analysis, data interpretation, and drafting of manuscript.

## Competing interests

The authors have filed a patent (U.S. 11,696,960) regarding the use of *N*-aminoanthranilic acids for aldehyde imaging.

## Data and materials availability

All raw and source data are available on request of the corresponding author.

## Notes

### Competing Interest Statement

The authors have declared no competing interest.

## REFERENCES

[1] J. D. Cassidy, L. Carroll, P. Peloso, J. Borg, H. Von Holst, L. Holm, J. Kraus, V. Coronado, Journal of rehabilitation medicine 2004, 36, 28–60.

[2] G. T. Baldwin, M. J. Breiding, R. D. Comstock, Handbook of clinical neurology 2018, 158, 63–74.

[3] S. Stephenson, J. Langley, C. Cryer, Accident Analysis & Prevention 2005, 37, 825–832.

[4] aP. H. Montenigro, M. L. Alosco, B. M. Martin, D. H. Daneshvar, J. Mez, C. E. Chaisson, C. J. Nowinski, R. Au, A. C. McKee, R. C. Cantu, Journal of neurotrauma 2017, 34, 328–340; bA. Buki, N. Kovacs, E. Czeiter, K. Schmid, R. P. Berger, F. Kobeissy, D. Italiano, R. L. Hayes, F. C. Tortella, E. Mezosi, Advances and Technical Standards in Neurosurgery: Volume 42 2015, 147-192.

[5] H. F. Lingsma, M. C. Cnossen, The Lancet Neurology 2017, 16, 494–495.

[6] aM. J. Haydel, C. A. Preston, T. J. Mills, S. Luber, E. Blaudeau, P. M. DeBlieux, New England Journal of Medicine 2000, 343, 100–105; bC. J. Homer, L. Kleinman, Pediatrics 1999, 104, e78-e78.

[7] A. Olson, M. J. Ellis, E. Selci, K. Russell, Frontiers in neurology 2020, 11, 220.

[8] C. C. Giza, J. S. Kutcher, S. Ashwal, J. Barth, T. S. Getchius, G. A. Gioia, G. S. Gronseth, K. Guskiewicz, S. Mandel, G. Manley, Neurology 2013, 80, 2250–2257.

[9] C. C. Giza, D. A. Hovda, Neurosurgery 2014, 75, S24.

[10] aZ. Yu, W. Li, U. T. Brunk, Apmis 2003, 111, 643–652; bF. Petronilho, G. Feier, B. de Souza, C. Guglielmi, L. S. Constantino, R. Walz, J. Quevedo, F. Dal-Pizzol, Journal of Surgical Research 2010, 164, 316-320; cA. Halstrom, E. MacDonald, C. Neil, G. Arendts, D. Fatovich, M. Fitzgerald, ournal of Clinical Neuroscience 2017, 35, 104-108; dL. Cristofori, B. Tavazzi, R. Gambin, R. Vagnozzi, S. Signoretti, A. M. Amorini, G. Fazzina, G. Lazzarino, Clinical biochemistry 2005, 38, 97-100; eK. Zahedi, F. Huttinger, R. Morrison, T. Murray-Stewart, R. A. Casero Jr, K. I. Strauss, Journal of neurotrauma 2010, 27, 515-525; fN. J. Yates, S. Lydiard, B. Fehily, G. Weir, A. Chin, C. A. Bartlett, J. Alderson, M. Fitzgerald, Experimental brain research 2017, 235, 2133-2149.

[11] I. N. Singh, L. K. Gilmer, D. M. Miller, J. E. Cebak, J. A. Wang, E. D. Hall, Journal of cerebral blood flow & metabolism 2013, 33, 593–599.

[12] aJ. E. Cebak, I. N. Singh, R. L. Hill, J. A. Wang, E. D. Hall, Journal of neurotrauma 2017, 34, 1302–1317; bE. D. Hall, R. A. Vaishnav, A. G. Mustafa, Neurotherapeutics 2010, 7, 51-61; cC. Shao, K. N. Roberts, W. R. Markesbery, S. W. Scheff, M. A. Lovell, Free Radical Biology and Medicine 2006, 41, 77-85.

[13] H. Isokuortti, G. L. Iverson, A. Kataja, A. Brander, J. Öhman, T. M. Luoto, Journal of neurotrauma 2016, 33, 232–241.

[14] R. S. Sohal, W. C. Orr, Free Radical Biology and Medicine 2012, 52, 539–555.

[15] aH. Esterbauer, R. J. Schaur, H. Zollner, Free radical Biology and medicine 1991, 11, 81–128; bA. Ayala, M. F. Muñoz, S. Argüelles, Oxidative medicine and cellular longevity 2014, 2014.

[16] B. Roozenbeek, A. I. Maas, D. K. Menon, Nature Reviews Neurology 2013, 9, 231–236.

[17] C. H. Hume, B. J. Wright, G. J. Kinsella, Journal of the International Neuropsychological Society 2022, 28, 736–755.

[18] J. Chen, B. Yu, Aging Clinical and Experimental Research 1996, 8, 334–340.

[19] aE. J. Anderson, L. A. Katunga, M. S. Willis, Clinical and Experimental Pharmacology and Physiology 2012, 39, 179–193; bB. Yoval-Sánchez, J. S. Rodríguez-Zavala, Chemical research in toxicology 2012, 25, 722-729.

[20] K. Kitagawa, T. Kawamoto, N. Kunugita, T. Tsukiyama, K. Okamoto, A. Yoshida, K. Nakayama, K.-i. Nakayama, FEBS letters 2000, 476, 306–311.

[21] T. Isse, T. Oyama, K. Kitagawa, K. Matsuno, A. Matsumoto, A. Yoshida, K. Nakayama, K.-i. Nakayama, T. Kawamoto, Pharmacogenetics and Genomics 2002, 12, 621–626.

[22] A. Loan, J. W.-H. Leung, D. P. Cook, C. Ko, B. C. Vanderhyden, J. Wang, H. M. Chan, Iscience 2023, 26.

[23] H. Azari, S. Sharififar, M. Rahman, S. Ansari, B. A. Reynolds, J Vis Exp 2011, 47.

[24] aE. Shohami, M. Novikov, R. Bass, Brain research 1995, 674, 55–62; bL. Beni-Adani, I. Gozes, Y. Cohen, Y. Assaf, R. A. Steingart, D. E. Brenneman, O. Eizenberg, V. Trembolver, E. Shohami, Journal of Pharmacology and Experimental Therapeutics 2001, 296, 57-63; cM. A. Flierl, P. F. Stahel, K. M. Beauchamp, S. J. Morgan, W. R. Smith, E. Shohami, Nature protocols 2009, 4, 1328-1337; dI. Khalin, N. L. A. Jamari, N. B. A. Razak, Z. B. Hasain, M. A. bin Mohd Nor, M. H. bin Ahmad Zainudin, A. B. Omar, R. Alyautdin, Neural regeneration research 2016, 11, 630.

[25] O. Chao, M. Pum, J.-S. Li, J. Huston, Neuroscience 2012, 202, 318–325.

[26] M. Suchý, C. Lazurko, A. Kirby, T. Dang, G. Liu, A. J. Shuhendler, Org Biomol Chem 2019, 17, 1843–1853.

[27] A. Kirby, M. Suchý, A. Brouwer, A. Shuhendler, Chemical Communications 2019, 55, 5371–5374.

[28] C. S. McKay, M. Finn, Angewandte Chemie 2016, 128, 12833–12839.

[29] I. Blockx, S. Einstein, P.-J. Guns, J. Van Audekerke, C. Guglielmetti, W. Zago, D. Roose, M. Verhoye, A. Van der Linden, F. Bard, Journal of Alzheimer’s Disease 2016, 54, 723–735.

[30] G. Liu, X. Song, K. W. Chan, M. T. McMahon, NMR in Biomedicine 2013, 26, 810–828.

[31] J. C. Er, C. Leong, C. L. Teoh, Q. Yuan, P. Merchant, M. Dunn, D. Sulzer, D. Sames, A. Bhinge, D. Kim, Angewandte Chemie International Edition 2015, 54, 2442–2446.

[32] M. T. Scimone, H. C. Cramer III, P. Hopkins, J. B. Estrada, C. Franck, Plos one 2020, 15, e0229520.

[33] aY. Xiong, A. Mahmood, M. Chopp, Nature Reviews Neuroscience 2013, 14, 128–142; bA. L. Petraglia, M. L. Dashnaw, R. C. Turner, J. E. Bailes, Neurosurgery 2014, 75, S34-S49.

[34] D. F. Meaney, D. H. Smith, Clinics in sports medicine 2011, 30, 19–31.

[35] aK. D. Statler, H. Alexander, V. Vagni, R. Holubkov, C. E. Dixon, R. S. Clark, L. Jenkins, P. M. Kochanek, Brain research 2006, 1076, 216–224; bK. D. Statler, H. Alexander, V. Vagni, C. E. Dixon, R. S. Clark, L. Jenkins, P. M. Kochanek, Journal of neurotrauma 2006, 23, 97-108.

[36] aM. Makdissi, D. Darby, P. Maruff, A. Ugoni, P. Brukner, P. R. McCrory, The American journal of sports medicine 2010, 38, 464–471; bM. P. McClincy, M. R. Lovell, J. Pardini, M. W. Collins, M. K. Spore, Brain injury 2006, 20, 33-39; cJ. Ponsford, C. Willmott, A. Rothwell, P. Cameron, A.-M. Kelly, R. Nelms, C. Curran, K. Ng, Journal of the International Neuropsychological Society 2000, 6, 568-579.

[37] aA. Meconi, R. C. Wortman, D. K. Wright, K. J. Neale, M. Clarkson, S. R. Shultz, B. R. Christie, PloS one 2018, 13, e0197187; bA. L. Petraglia, B. A. Plog, S. Dayawansa, M. Chen, M. L. Dashnaw, K. Czerniecka, C. T. Walker, T. Viterise, O. Hyrien, J. J. Iliff, Journal of neurotrauma 2014, 31, 1211-1224.

[38] R. D. Brady, P. M. Casillas-Espinosa, D. V. Agoston, E. H. Bertram, A. Kamnaksh, B. D. Semple, S. R. Shultz, Neurobiology of disease 2019, 123, 8–19.

[39] Y. K. Baskin, W. D. Dietrich, E. J. Green, Journal of neuroscience methods 2003, 129, 87–93.

[40] aA. K. Heye, R. D. Culling, M. d. C. V. Hernández, M. J. Thrippleton, J. M. Wardlaw, NeuroImage: Clinical 2014, 6, 262–274; bL. C. Turtzo, N. Jikaria, M. R. Cota, J. P. Williford, V. Uche, T. Davis, J. MacLaren, A. D. Moses, G. Parikh, M. A. Castro, Brain Communications 2020, 2, fcaa143.

[41] N. Patel, O. Kirmi, in Seminars in Ultrasound, CT and MRI, Vol. 30, Elsevier, 2009, pp. 559–564.

[42] T. Dang, M. Suchý, Y. J. Truong, W. Oakden, W. W. Lam, C. Lazurko, G. Facey, G. J. Stanisz, A. J. Shuhendler, Chemistry – A European Journal 2018, 24, 9148–9156.

[43] E. M. S. P.-T. Brun, N. D. Calvert, M. Suchý, A. Kirby, G. Melkus, R. Garipov, C. L. Addison, A. J. Shuhendler, Chemical Communications 2021, 57, 10867–10870.

[44] A. Guillot, T. Ren, T. Jourdan, R. J. Pawlosky, E. Han, S.-J. Kim, L. Zhang, G. F. Koob, B. Gao, Proceedings of the National Academy of Sciences 2019, 116, 25974–25981.

[45] C. Farina, F. Aloisi, E. Meinl, Trends in Immunology 2007, 28, 138–145.

[46] aK. Walus, S. Beyer, S. M. Willerth, Current Opinion in Biomedical Engineering 2020, 14, 25–33; bM. D. Tang-Schomer, J. D. White, L. W. Tien, L. I. Schmitt, T. M. Valentin, D. J. Graziano, A. M. Hopkins, F. G. Omenetto, P. G. Haydon, D. L. Kaplan, Proceedings of the National Academy of Sciences 2014, 111, 13811-13816; cD. Warren, E. Tomaskovic-Crook, G. G. Wallace, J. M. Crook, APL bioengineering 2021, 5.

[47] aW. Lee, J. Pinckney, V. Lee, J.-H. Lee, K. Fischer, S. Polio, J.-K. Park, S.-S. Yoo, Neuroreport 2009, 20, 798–803; bD. N. Rockwood, R. C. Preda, T. Yücel, X. Wang, M. L. Lovett, D. L. Kaplan, Nature protocols 2011, 6, 1612-1631; cR. Nazarov, H.-J. Jin, D. L. Kaplan, Biomacromolecules 2004, 5, 718-726.

[48] D. Bouvier, C. Oris, M. Brailova, J. Durif, V. Sapin, Clinical biochemistry 2020, 85, 5–11.

[49] J. J. Bazarian, P. Biberthaler, R. D. Welch, L. M. Lewis, P. Barzo, V. Bogner-Flatz, P. G. Brolinson, A. Büki, J. Y. Chen, R. H. Christenson, The Lancet Neurology 2018, 17, 782–789.

[50] aS. Çevik, M. M. Özgenç, A. Güneyk, Ş. Evran, E. Akkaya, F. ÇaliŞ, S. Katar, C. Soyalp, H. Hanımoğlu, M. Y. Kaynar, Clinical neurology and neurosurgery 2019, 183, 105380; bN. A. Huebschmann, T. M. Luoto, J. E. Karr, K. Berghem, K. Blennow, H. Zetterberg, N. J. Ashton, J. Simrén, J. P. Posti, J. M. Gill, Frontiers in Neurology 2020, 11, 1054; cE. Czeiter, K. Amrein, B. Y. Gravesteijn, F. Lecky, D. K. Menon, S. Mondello, V. F. Newcombe, S. Richter, E. W. Steyerberg, T. V. Vyvere, EBioMedicine 2020, 56.

[51] aJ. Gill, L. Latour, R. Diaz-Arrastia, V. Motamedi, C. Turtzo, P. Shahim, S. Mondello, C. DeVoto, E. Veras, D. Hanlon, Neurology 2018, 91, e1385–e1389; bM. Y. Mahan, M. Thorpe, A. Ahmadi, T. Abdallah, H. Casey, D. Sturtevant, S. Judge-Yoakam, C. Hoover, D. Rafter, J. Miner, World neurosurgery 2019, 128, e434-e444.

[52] N. R. Nichols, J. R. Day, N. J. Laping, S. A. Johnson, C. E. Finch, Neurobiology of aging 1993, 14, 421–429.

[53] aR. C. Gardner, R. Rubenstein, K. K. Wang, F. K. Korley, J. K. Yue, E. L. Yuh, P. Mukherje, A. B. Valadka, D. O. Okonkwo, R. Diaz-Arrastia, Journal of neurotrauma 2018, 35, 2341–2350; bO. Calcagnile, A. Holmén, M. Chew, J. Undén, Scandinavian journal of trauma, resuscitation and emergency medicine 2013, 21, 1-6.

[54] aP. K. Dash, J. Zhao, G. Hergenroeder, A. N. Moore, Neurotherapeutics 2010, 7, 100–114; bS. H. Chou, C. S. Robertson, P. i. t. I. M.-d. C. C. o. t. M. Monitoring, Neurocritical care 2014, 21, 187-214.

[55] P. J. Brooks, M.-A. Enoch, D. Goldman, T.-K. Li, A. Yokoyama, PLoS medicine 2009, 6, e1000050.

[56] B. Wu, L. Yu, Y. Wang, H. Wang, C. Li, Y. Yin, J. Yang, Z. Wang, Q. Zheng, H. Ma, Oncotarget 2016, 7, 2175.

[57] A. Kasai, E. Jee, Y. Tamura, K. Kouzaki, T. Kotani, K. Nakazato, American Journal of Physiology-Regulatory, Integrative and Comparative Physiology 2022, 322, R511–R525.

[58] aM. Comporti, Laboratory investigation; a journal of technical methods and pathology 1985, 53, 599–623; bP. A. Baeuerle, D. Baltimore, Cell 1996, 87, 13-20.

[59] M. Grossmann, Y. Nakamura, R. Grumont, S. Gerondakis, The international journal of biochemistry & cell biology 1999, 31, 1209–1219.

[60] aD. P. Clark, V. M. Perreau, S. R. Shultz, R. D. Brady, E. Lei, S. Dixit, J. M. Taylor, P. M. Beart, W. C. Boon, Neurochemical research 2019, 44, 1410–1424; bM. A. Wheeler, F. J. Quintana, Cold Spring Harbor perspectives in medicine 2019, 9; cE. C. Dresselhaus, M. K. Meffert, Frontiers in immunology 2019, 10, 1043.

[61] aM. Nonaka, X.-H. Chen, J. E. Pierce, M. J. Leoni, T. K. McIntosh, J. A. Wolf, D. H. Smith, Journal of neurotrauma 1999, 16, 1023–1034; bO. Sanz, L. Acarin, B. González, B. Castellano, Journal of neuroscience research 2002, 67, 772-780.

[62] L. Sun, M. Li, X. Ma, H. Feng, J. Song, C. Lv, Y. He, Journal of neuroinflammation 2017, 14, 1–18.

[63] C. Pan, J.-h. Xing, C. Zhang, Y.-m. Zhang, L.-t. Zhang, S.-j. Wei, M.-x. Zhang, X.-p. Wang, Q.-h. Yuan, L. Xue, Oncotarget 2016, 7, 35562.

[64] R. Saggu, T. Schumacher, F. Gerich, C. Rakers, K. Tai, A. Delekate, G. C. Petzold, Acta neuropathologica communications 2016, 4, 1–10.

[65] aS. Dwivedi, A. Sharma, B. Patrick, R. Sharma, Y. C. Awasthi, Redox Report 2007, 12, 4–10; bJ. M. Müller, R. A. Rupec, P. A. Baeuerle, Methods 1997, 11, 301-312; cM. Meyer, H. L. Pahl, P. A. Baeuerle, Chemico-biological interactions 1994, 91, 91-100.

[66] aU. C. Yadav, K. V. Ramana, Y. C. Awasthi, S. K. Srivastava, Toxicology and applied pharmacology 2008, 227, 257–264; bN. Žarković, K. Žarković, R. J. Schaur, S. Štolc, G. Schlag, H. Redl, G. Waeg, S. Borović, I. Lončarić, G. Jurić, Life Sciences 1999, 65, 1901-1904; cC. A. Thompson, P. C. Burcham, Chemical Research in Toxicology 2008, 21, 2245-2256.

[67] N. Kikuchi, T. Tajima, Y. Tamura, Y. Yamanaka, K. Menuki, T. Okamoto, M. Sakamaki-Sunaga, A. Sakai, K. Hiranuma, K. Nakazato, Biology of Sport 2022, 39, 429–434.

[68] aL. M. Kenny, E. O. Aboagye, Advances in cancer research 2014, 124, 329–374; bR. C. Shaw, G. D. Tamagnan, A. A. S. Tavares, Frontiers in Neuroscience 2020, 14, 871; cG. C. Van de Bittner, E. L. Ricq, J. M. Hooker, Accounts of chemical research 2014, 47, 3127-3134.

