## Supplemental material for "Aldehydic load as an objective imaging biomarker of mild traumatic brain injury"

**Supplemental figures and table**

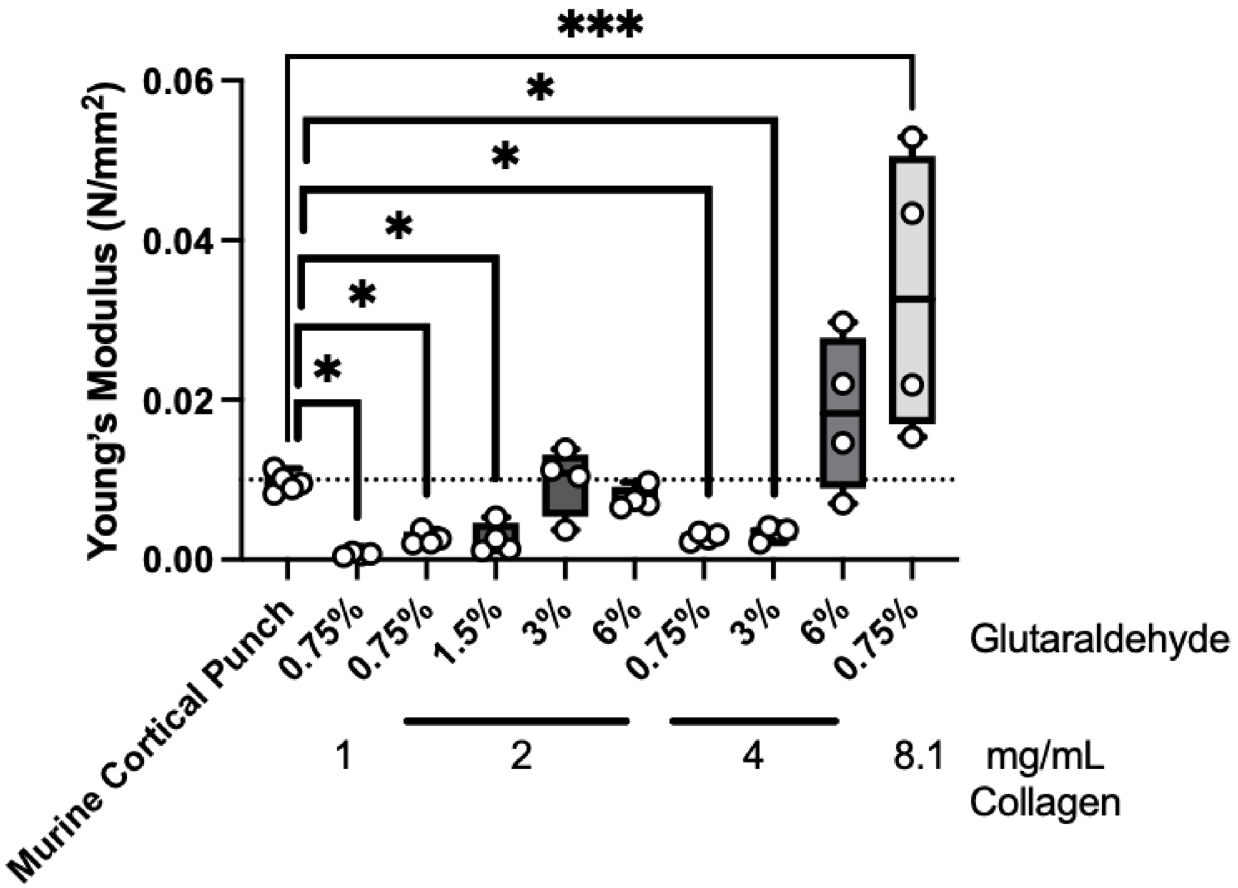


**Fig. S1. Young's Modulus of different formulations of corticomimetic scaffold exterior in comparison to murine cortical punch**. Various concentrations of glutaraldehyde cross-linker and collagen (n=4 per group) were tested and compared to a punch of cortical brain tissue of a mouse (n=5) using a universal testing machine (UTM). Boxplots and individual data points are shown, *p<0.05.

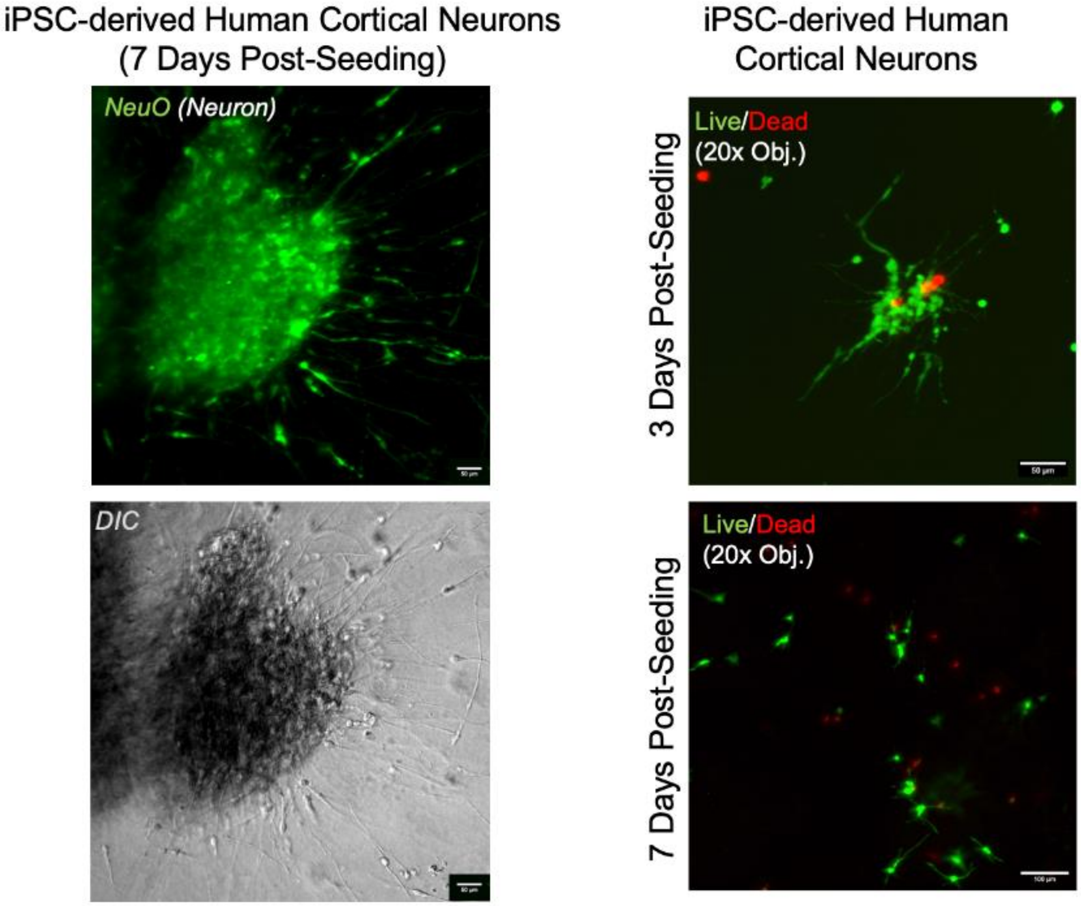


**Fig. S2. 3D corticomimetic scaffold made for *in vitro* concussion model can support growth of primary human neurons.** iPSC-derived human cortical neurons were seeded into the central neural plug of the corticomimetic scaffold. NeuO staining was performed and imaged on a confocal microscope 7 days post-seeding to confirm differentiation of neurons (left; scale bar= 50µM). A majority of cells were alive following live/dead staining with calcein-AM (green; alive) and ethidium homodimer-1 (red/dead) at 3 days- (top) and 7-days (bottom) post-seeding. Scale = 100 µM.

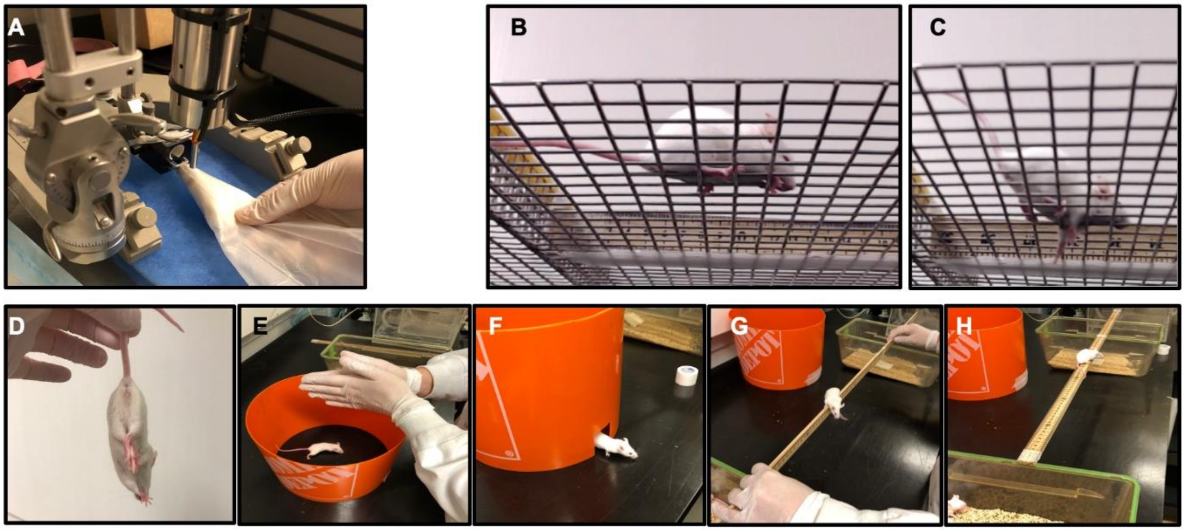


**Fig. S3. Awake closed head impact (ACHI) setup and behavioural tests for mouse model of mTBI.** (**A**) Awake mice are restrained in a conical bag with an opening to allow for breathing. The mouse is then placed under a 5mm impactor tip held by a stereotaxic rig, with filter floss placed underneath to allow free rotation of the head and neck. (**B-C**) Examples of mice performing the grid walk test where they are allowed to walk around a raised grid for three minutes before (B) and after (C) ACHI, where foot faults and activity time are recorded. (**D-H**) Five pass-or-fail tests are conducted before and after ACHI which comprise the neurological severity score (NSS). (D) Mice are hung from the tail and awarded a point (fail) if the hindlimbs cross; (E) startle reflex, where the researcher claps and a point is awarded for any reaction to the sound; (F) escape paradigm, where mice are placed in the middle of a ~25cm diameter cylinder with a door and given 3 minutes to escape; (G) balance test, where mice are hung by the front paws and given 10 seconds to lift themselves into a perched position as in the photo; (H) beam walk test where a mouse is placed on a meter stick 30cm away from a cage (one home cage, one new cage) on either end and given 3 minutes to reach a cage.

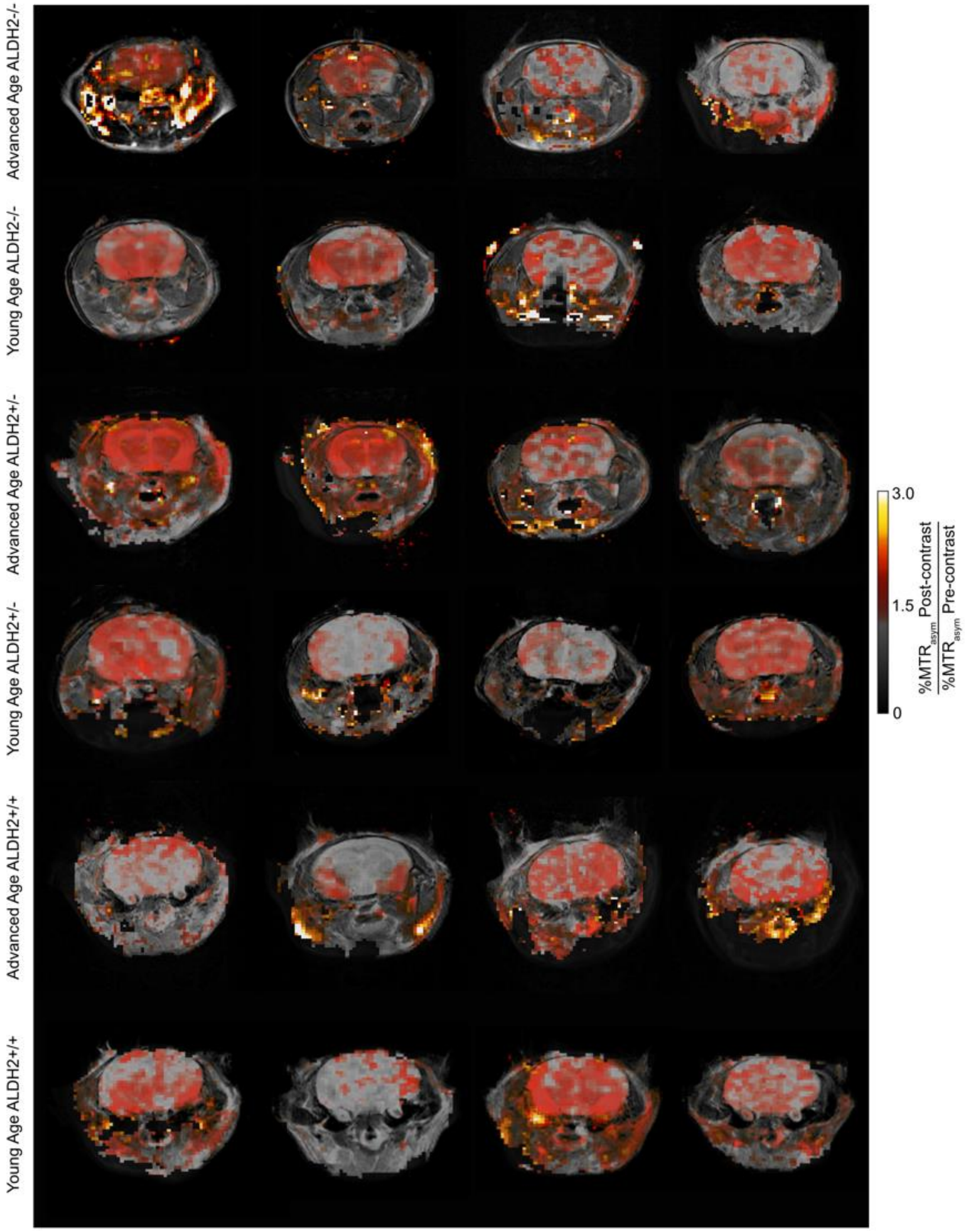


**Fig. S4. Mapping of brain aldehydes by ProxyNA$_3$ in mice two days post-mTBI.** Mice separated into groups by age (young <16 weeks, aged >65 weeks) and genotype (ALDH2 $^{+/+}$, $^{+/-}$, $^{-/-}$) underwent CEST-MRI at two days post-concussive impact. %MTR$_{asym}$ after ProxyNA$_3$ injection normalized to pre-contrast signal is mapped onto coronal T$_2$-weighted image for each mouse. There is sufficient binding of ProxyNA$_3$ to endogenous aldehydes for detection in every mouse two days post-mTBI.

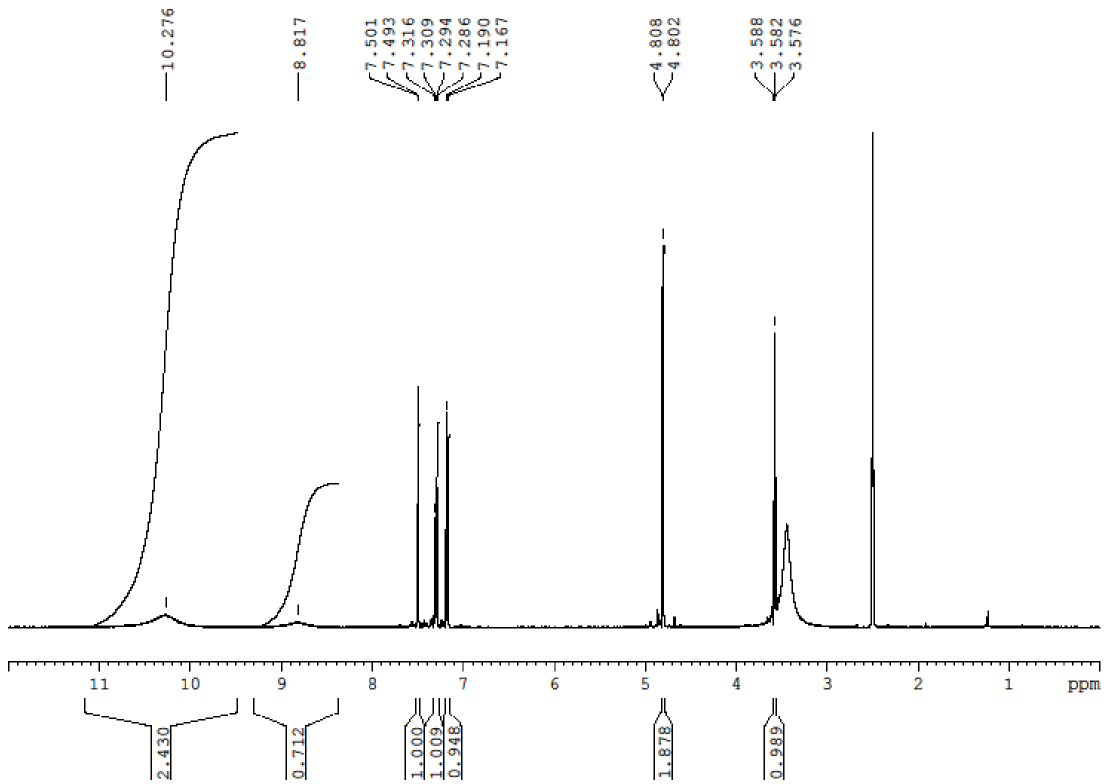


**Fig. S5** $^{1}$H NMR spectrum of ProxyNA$_3$. Signal at δ 3.56 ppm indicates the presence of small amount of dioxane in the sample.

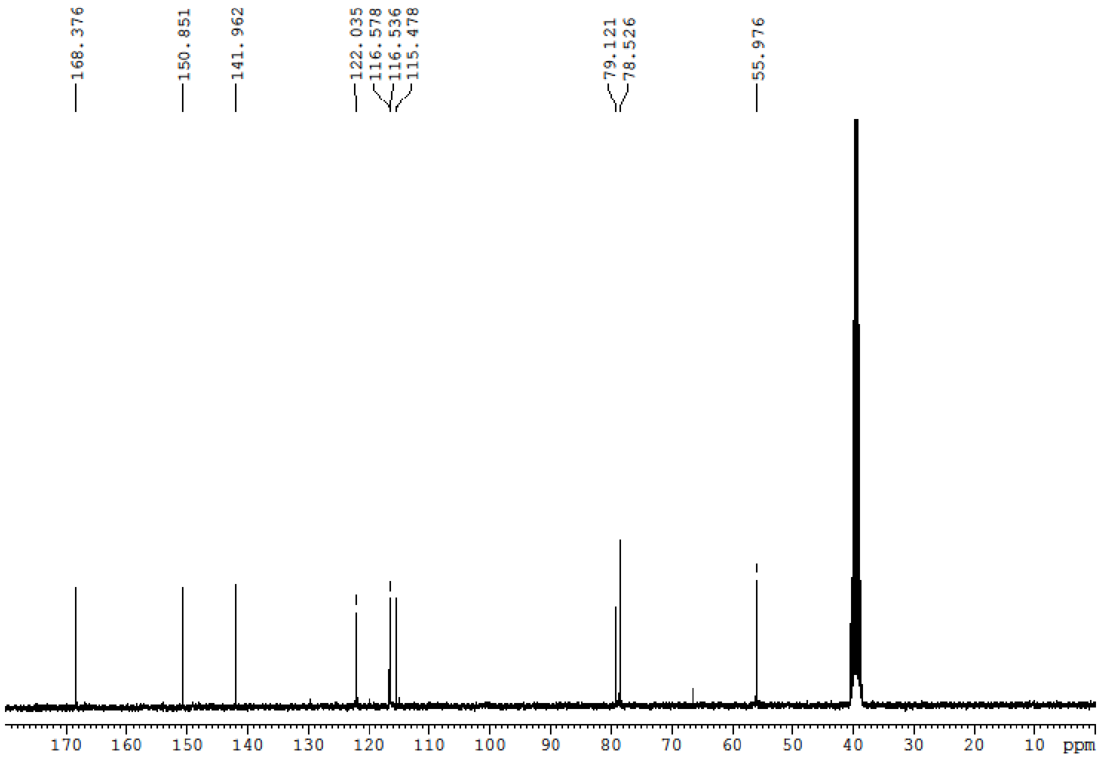


**Fig. S6** $^{13}C$ NMR spectrum of ProxyNA$_3$.

**Table S1. Impact velocities tested for mouse model of mTBI using electromechanical impactor device.** Three impact velocities were evaluated using a 5mm tip on an impactor device based on previously reported models of mTBI. Behavioural deficits were only noted for 5 and 6m/s impacts. Less than 5% of mice impacted at 5m/s elicited an unwanted outcome, compared to 100% impacted at 6m/s.

| Impact Velocity | Behavioural outcome | Unwanted outcome* | References |
|---|---|---|---|
| 2.35m/s | None | None | [1] |
| 5m/s | Present | <5% | [2] |
| 6m/s | Present | 100% | [3]** |

*Unwanted outcome= severe TBI (seizure, skull fracture, brain hemorrhaging, death)
**6.8m/s; our impactor device had maximum velocity of 6.0m/s

**Movie S1. Grid walk recording of wild type mouse pre-mTBI.** Mice are placed on a raised grid and allowed to walk freely for 3 minutes. Total number of forelimb steps, foot faults, and time spent active is counted by a blinded researcher. The naïve mouse walks across the grid with no foot faults.

**Movie S2. Grid walk recording of wild type mouse post-mTBI.** Mice are placed on a raised grid and allowed to walk freely for 3 minutes. Total number of forelimb steps, foot faults, and time spent active is counted by a blinded researcher. The video shows a mouse post-concussive impact walking across the grid with many foot faults.

**Supplementary References**


[1] aB. S. Main, S. S. Sloley, S. Villapol, D. N. Zapple, M. P. Burns, *JoVE (Journal of Visualized Experiments)* **2017**, e55713; bC. N. Winston, A. Noël, A. Neustadtl, M. Parsadanian, D. J. Barton, D. Chellappa, T. E. Wilkins, A. D. Alikhani, D. N. Zapple, S. Villapol, *The American journal of pathology* **2016**, *186*, 552-567.

[2] aA. D. Bachstetter, S. J. Webster, D. S. Goulding, J. E. Morton, D. M. Watterson, L. J. Van Eldik, *Journal of neuroinflammation* **2015**, *12*, 1-9; bJ. A. Creed, A. M. DiLeonardi, D. P. Fox, A. R. Tessler, R. Raghupathi, *J Neurotrauma* **2011**, *28*, 547-563; cM. T. Goodus, N. A. Kerr, R. Talwar, D. Buziashvili, J. E. Fragale, K. C. Pang, S. W. Levison, *Journal of Neurotrauma* **2016**, *33*, 1522-1534; dB. L. Schneider, F. Ghoddoussi, J. L. Charlton, R. J. Kohler, M. P. Galloway, S. A. Perrine, A. C. Conti, *J Neurotrauma* **2016**, *33*, 1614-1624; eS. J. Webster, L. J. Van Eldik, D. M. Watterson, A. D. Bachstetter, *The Journal of neuroscience : the official journal of the Society for Neuroscience* **2015**, *35*, 6554-6569; fK. M. Wendel, J. B. Lee, B. M. Affeldt, M. Hamer, I. S. Harahap-Carrillo, A. C. Pardo, A. Obenaus, *J Neurotrauma* **2018**; gT. Emmerich, L. Abdullah, J. Ojo, B. Mouzon, T. Nguyen, G. Crynen, J. E. Evans, J. Reed, M. Mullan, F. Crawford, *Neuromolecular medicine* **2017**, *19*, 122-135; hX. Han, Z. Chai, X. Ping, L.-J. Song, C. Ma, Y. Ruan, X. Jin, *Frontiers in neuroscience* **2020**, *14*, 210; iY. Shitaka, H. T. Tran, R. E. Bennett, L. Sanchez, M. A. Levy, K. Dikranian, D. L. Brody, *Journal of Neuropathology & Experimental Neurology* **2011**, *70*, 551-567; jA. G. Velosky, L. B. Tucker, A. H. Fu, J. Liu, J. T. McCabe, *Behavioural brain research* **2017**, *324*, 115-124; kB. Mouzon, H. Chaytow, G. Crynen, C. Bachmeier, J. Stewart, M. Mullan, W. Stewart, F. Crawford, *Journal of neurotrauma* **2012**, *29*, 2761-2773.

[3] J. R. Lynch, H. Wang, B. Mace, S. Leinenweber, D. S. Warner, E. R. Bennett, M. P. Vitek, S. McKenna, D. T. Laskowitz, *Experimental neurology* **2005**, *192*, 109-116.
